# Beta-Adrenergic Stimulation and *MYH7* G256E Mutant Gene Dosage Drive Hypertrophic Cardiomyopathy Phenotype Penetrance

**DOI:** 10.64898/2026.06.02.729411

**Authors:** Paul Heinrich, Raina M. Jung, Jonathan S. Achter, Vincent X. Nguyen, Carissa A. Lee, Carolin Sailer, Henna Domian, Alison Vander Roest, Fabian P. Suchy, James W. Jahng, Ana Kojic, Daniel Lee, Amelie Paasche, Brock Roberts, Hiromitsu Nakauchi, Han Zhu, Joseph C. Wu, Daniel Bernstein, Alessandra Moretti, Alicia Lundby, Soah Lee, Sean M. Wu

## Abstract

**Aims:** Hypertrophic cardiomyopathy (HCM) is the most prevalent genetic heart disorder, characterized by significant phenotypic variability even among individuals with identical *MYH7* mutations. This study aims to elucidate factors contributing to this variability and identify drivers of phenotype penetrance. We compared the baseline phenotypes of a highly penetrant *MYH7* H251N mutation and the variably penetrant *MYH7* G256E mutation and investigated the impact of adding beta-adrenergic stimulation and homozygosity on disease phenotype penetrance using cardiomyocytes from an isogenic line of human induced pluripotent stem cells (hiPSC-CMs).

**Methods and Results:** Isogenic hiPSCs with *MYH7* H251N and *MYH7* G256E mutations were generated using CRISPR/Cas9 technology and differentiated into cardiomyocytes (CMs). Single-cell RNA sequencing (scRNAseq) and functional analysis of contractile function revealed consistent HCM phenotype presentation in H251N CMs, whereas G256E CMs exhibited a subtle and more variable phenotype. Beta-adrenergic stimulation induced a distinct metabolic stress response in G256E CMs, characterized by impaired mitochondrial ATP upregulation. Increasing mutant gene dosage from hetero- to homozygosity led to consistent increase in hypertrophic and structural gene expression changes in G256E CMs at RNA and protein levels. These changes were distinct from the changes observed with stress response. Importantly, homozygous G256E CMs exhibited a hypercontractile functional and disorganized structural phenotype. Across multiple experimental conditions, we identified consistent increase in cardiomyocyte specific transcriptomic markers such as *NPPB*, *APOE*, *PDLIM3* and *ANKRD1*.

**Conclusions:** Our study highlights the use of a variably penetrant MYH7 mutation to investigate factors that influence HCM phenotype penetrance. Specifically, we found that mutant gene dosage and beta-adrenergic stimulation induce distinct HCM disease phenotypes, providing novel insights into mechanisms that may contribute to variable disease expression in HCM.

**Translational Perspective:** HCM is characterized by significant phenotypic variability, complicating both diagnosis and clinical management. This study explores the factors driving HCM phenotype penetrance using isogenic hiPSC-CMs with *MYH7* mutations. We demonstrate that beta-adrenergic stimulation and increased mutant gene dosage significantly impact HCM disease penetrance. Beta-adrenergic stimulation triggers metabolic stress responses, while increased gene dosage leads to a hypercontractile and structurally disorganized phenotype. These findings provide insight into how specific modifiers can shape disease-associated phenotypes in HCM model systems.

**Graphical Abstract:** 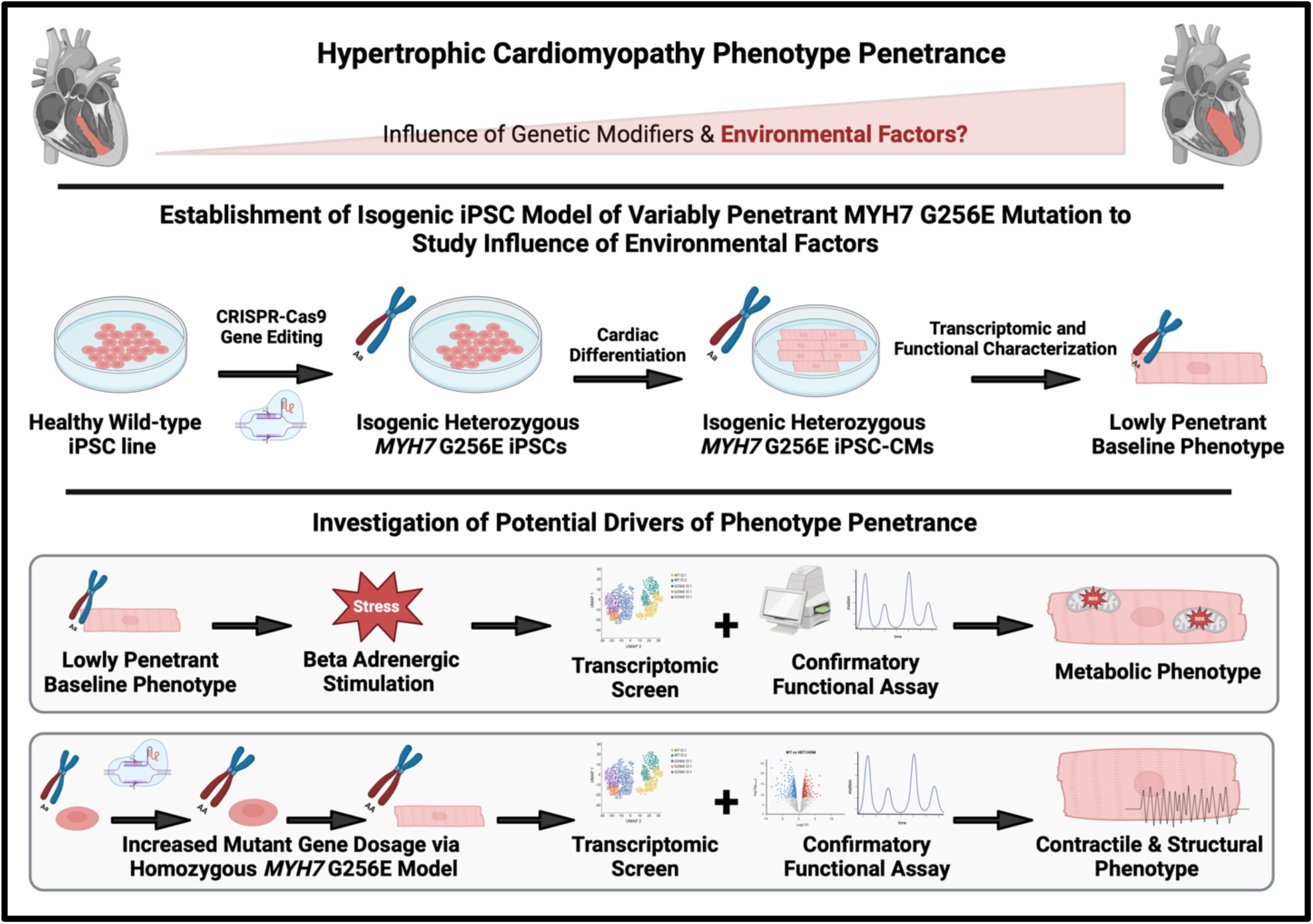

## 1. Introduction

HCM is the most prevalent genetic heart disorder, affecting up to 1 in 200 individuals. It is primarily characterized by cardiac hypercontractility, hypertrophy, myocyte and myofibril disarray, and interstitial fibrosis. Complications of HCM often include left ventricular outflow obstruction, heart failure, arrhythmia, and sudden cardiac death.^1^ Approximately 60% of HCM cases are attributed to the inheritance of a single autosomal dominant mutation in genes coding for sarcomere proteins. Among these, the *MYH7* gene, which encodes the β-myosin heavy chain (MHC), is particularly noteworthy, with over 200 mutations implicated in HCM accounting for around a third of all cases.^2^ Despite the high number of known mutations associated with HCM, the degree to which each mutation contributes to the HCM phenotype remains largely unknown.^3^ While some mutations are associated with an early onset of HCM during childhood such as *MYH7* H251N or *MYH7* P710R^4^, most mutations exhibit highly variable clinical penetrance. A recent metanalysis of 17 studies revealed that *MYH7* mutations have an average phenotype penetrance of 64.3%, with a 95% confidence interval of 52.6%-74.5%.^5^ An illustrative example of a clinically variable *MYH7* mutation is G256E. A larger kindred study of this mutation revealed that 19 out of 34 adults and 3 out of 5 children (1-11 years old) exhibited left ventricular hypertrophy, representing a wide range of phenotype penetrance, from very early onset in childhood to progressive HCM development throughout adulthood, or no hypertrophy phenotype development at all.^6^

The variability in phenotype penetrance among these cases is hypothesized to be influenced by both genetic and non-genetic modifiers. Genetic modifiers include inherited or de novo occurring single nucleotide polymorphisms (SNPs), which can increase the phenotype penetrance of a known mutation. By comparing hiPSC-CMs from siblings both carrying the same HCM-associated mutation with only one exhibiting a clinical HCM phenotype, Escriba et al. demonstrated the partial contribution of SNPs to phenotype penetrance by correcting a SNP present in a different sarcomere gene between the siblings. This led to a partial rescue of the HCM phenotype thereby supporting the partial contribution of the SNP to disease penetrance.^7^

Conversely, two independent monozygotic twin studies reported drastically differing clinical phenotypes between twins despite having identical genotypes.^8^ Repetti et al. reported a 5–14-year follow-up of 11 monozygotic twin pairs, all presenting with discordant morphologic features of the heart. Whole genome sequencing in six of these twins did not reveal somatic genetic variants as a possible explanation.^8^ These studies strongly suggest that non-heritable and environmental factors can significantly contribute to the variable disease penetrance in HCM.

One theory for a non-heritable mechanism of variable HCM disease penetrance is allelic balance. HCM-associated *MYH7* mutations are mostly heterozygous missense mutations, with varying contribution of each allele to the final protein abundance.^9,10,11^ Allelic imbalance can be caused by numerous factors such as differing activity of cis-regulatory elements on each allele^9,10^ or by transcriptional bursts.^11^ Consequently, this imbalance can lead to permanent or transient changes in abundance of the mutant protein, which is hypothesized to correlate with a variable HCM phenotype. The exact molecular and functional consequences of increased or decreased mutant gene dosage however remain unknown.

Another potential non-heritable contributor to variable penetrance of HCM is environmental factors. Influences such as hypertension and obesity have been described to negatively impact HCM development.^12,13^ Conversely, beta-blockers can improve symptoms and increase survival in early-onset HCM and are the mainstay of treatment for dilated cardiomyopathies, reducing sympathetic drive.^14^ While these correlations have been described in epidemiologic and clinical studies, a mechanistic understanding of how environmental factors influence HCM penetrance is lacking, thus representing an unexplored opportunity for identifying disease-modifying pathways and potential drug targets.

Since the discovery of hiPSCs and subsequent development of directed cardiac differentiation protocols,^15^ hiPSC-CMs have been extensively used in disease modeling and drug development.^16^ One strength of this system is the ability to model a patient’s disease by generating patient-derived cell lines.^17^ Another application is the investigation of specific mutation effects in isogenic cell lines from healthy donors by introducing only the mutation of interest without altering the genomic background.^18^ Together with functional studies of recombinant proteins, this approach has led to the mechanistic understanding of highly penetrant HCM mutations such as *MYH7* P710R or *MYH7* H251N^19,20^ and has more recently been used to investigate less penetrant mutations such as *MYH7* G256E.^18^

In this study, we further develop the hiPSC-CM model to examine the HCM phenotype and factors influencing its variability in hiPSCs carrying a variably penetrant mutation (*MYH7* G256E), using a highly penetrant mutation (*MYH7* H251N) as a positive benchmark. The early-onset H251N mutation, located in the central β-sheet of the transducer region, increases ATPase activity, actin-gliding velocity, and intrinsic force, consistent with enhanced cross-bridge cycling.^19,20^ In contrast, the nearby G256E mutation lies within a β-hairpin of the transducer region and perturbs structural communication between nucleotide- and actin-binding sites, leading to increased myosin head availability and a higher duty ratio despite reduced actin gliding velocity.^18^

Our central hypothesis is that stress factors and mutant gene dosage are key contributors to disease penetrance in HCM. Specifically, we focus on the response to beta-adrenergic stimulation induced by isoproterenol and the effect of increased gene dosage in the context of cell-to-cell allelic imbalance by comparing heterozygous (HET) and homozygous (HOM) mutant *MYH7* G256E lines. In addition, to address the variability inherent in the hiPSC system, we analyze multiple clonally derived isogenic cell lines across independent replicates for each condition to assess the effect of run-to-run and clone-to-clone variability on phenotype penetrance. Collectively, our studies illustrated the ability of beta-adrenergic stimulation and *MYH7* G256E mutant gene dosage to drive hypertrophic cardiomyopathy phenotype penetrance thus providing new insights on variably penetrant HCM mutations and opportunities for future discovery of novel drug targets.

## 2. Methods

Detailed information on Materials and Methods is available in the Supplementary material.

### 2.1 hiPSC Culture and CM Differentiation

The parental ACTN2-mEGFP cell line on WTC background and CRISPR targeted *MYH7* G256E wild type (WT) & heterozygous (HET) and *MYH7* H251N WT & HET cell lines were developed at the Allen Institute for Cell Science and are available through the Coriell Institute Repository (https://www.coriell.org). Homozygous (HOM) *MYH7* G256E lines were generated by re-targeting the *MYH7* G256E HET line with the same mutant construct and are available on request.

hiPSCs were maintained in a 2D culture system coated with 1:400 Matrigel in DMEM/F12. Each hiPSC cell line was maintained in MTeSR hiPSC maintenance media. hiPSCs were differentiated into CMs using the canonical Wnt stimulation and inhibition protocol in RPMI 1640-based differentiation media according to Lian et al.^21^ On day 12, wells with over 90% beating CMs were treated with TrypLE Select Enzyme, centrifuged, resuspended in BamBanker freezing media, aliquoted into cryogenic vials, and stored at -80°C overnight before being transferred to a liquid nitrogen tank for long-term storage. For downstream experiments, frozen CMs were thawed, centrifuged, and plated in replating media (RPMI 1640 + B27 with 10% Knock Out Serum Replacement and Thiazovivin 1.0 μM). RPMI 1640 with B27 with insulin was used for CM maintenance.

### 2.2 Droplet-Based Single Cell RNA Sequencing

10X Genomics CellPlex technology was used to individually label CMs by underlying genotype and cell clone. Day 30 CMs were dissociated and resuspended in PBS with 0.04% BSA, labeled with Cell Multiplexing Oligo, washed, counted, pooled, and processed using the Chromium Controller Instrument. Libraries were sequenced on an Illumina NovaSeq instrument. Datasets were bioinformatically integrated and are publicly available under GEO accession number GSE276210.

### 2.3 scRNAseq Data Pre-Processing and Analysis

10X Cell Ranger Multi v7.2.0 was used to pre-process scRNAseq data, including sample demultiplexing, alignment, filtering, barcode processing, and gene counting. Reads were mapped to the human reference genome (GRCh38 2020-A). For downstream analysis, the Seurat R package was used.^22^ Cells with <200 or >10,000 detected genes and <250 or >150,000 RNA counts were filtered. Cells with >50% mitochondrial genes were also filtered. Raw read counts were normalized using the “LogNormalize” method. Non-linear dimensional reduction analysis such as UMAP and graph-based clustering were performed using Seurat. Non-myocyte clusters were removed based on *TNNT2* expression. The Harmony method was used to integrate different scRNAseq batches.^23^ Differentially expressed genes (DEGs) were identified, and gene ontology (GO) analysis was performed using the DAVID tool.

### 2.4 Contractility Measurement and Structural Assessment

Contraction velocity, defined as the maximal rate of displacement during the shortening phase of the contraction cycle, was measured in single hiPSC-CMs at day 35 of differentiation using the Sony SI8000 system. Cells were replated on day 27 at a density of approximately 40,000 cells/cm². On day 35, spontaneously beating individual CMs were recorded at 150 frames per second, and contraction velocity was quantified using edge-detection–based analysis implemented in the Sony SI8000 software.

For structural analysis, CMs were replated under the same conditions on day 27 and imaged on day 35. Z-disks were visualized using an α-actinin-GFP reporter and acquired on a Zeiss LSM980. Sarcomere orientation was quantified using OrientationJ (ImageJ), and fractional sarcomere shortening was analyzed using a modified version of SarcTrack implemented in MATLAB.^24^

### 2.5 Beta-adrenergic Stimulation & Analysis of Cell Metabolism (Seahorse)

Day 27 CMs were plated at 40,000 cells/cm^2^ for downstream contractility measurement, 50,000 cells/cm^2^ for downstream ATP rate assay and 160.000 cells/cm^2^ for downstream scRNAseq. Starting on day 30, CMs were treated with 1μM isoproterenol in RPMI 1640 supplemented with B27 with insulin for five days with media changes every day. Seahorse ATP Rate Assay was performed following the manufacturer’s instructions on day 35. OCR and ECAR were measured after treating hiPSC-CMs with oligomycin and rotenone/antimycin A.

### 2.6 Global Proteome Analysis & Data Processing

Day 30 hiPSC-CMs were lysed for proteome analysis. Three biological replicates were analyzed per clone. Protein concentration was determined, and samples were reduced, alkylated, digested, and desalted.^25^ Peptides were labeled with TMTpro reagents,^26^ fractionated, and analyzed on an Orbitrap Ascend Tribrid mass spectrometer. Raw files were analyzed with FragPipe v21.1. TMT16 workflow was used with default settings. Quantification results were filtered, and intensities were log2 transformed. Data are available via ProteomeXchange with identifier PXD054863.^27^ Differential protein abundance analysis was performed using a linear mixed model, using the ‘dream’ function from the varianceParition package.^28^ Functional enrichment was performed using the DAVID tool.

### 2.7 Statistical analysis

Data are presented as mean ± SEM unless otherwise indicated. Functional parameters were analyzed using linear mixed-effects models (lmerTest) with genotype/condition as a fixed effect and clone as a random effect; models were fit using maximum likelihood (REML = FALSE), and significance was assessed using Satterthwaite’s approximation. Comparisons between two groups were performed using unpaired two-tailed t tests, and comparisons among multiple groups were performed using ordinary one-way ANOVA with Tukey’s multiple-comparisons test. Differential gene expression in scRNA-seq data was assessed using the two-sided Wilcoxon rank-sum test with Bonferroni correction as implemented in Seurat. Proteomics P values were calculated using moderated t statistics with empirical Bayes moderation of standard errors, as implemented in limma, and adjusted for multiple testing using the Benjamini–Hochberg procedure. A value of P < 0.05 was considered statistically significant.

## 3. Results

### 3.1 Distinct Phenotype Penetrance & Variability of Different *MYH7* Mutations Is Reflected in hiPSC-CM Model

To examine whether the contrasting clinical phenotype penetrance of *MYH7* H251N and G256E mutations is reflected in the hiPSC model, three clonally derived isogenic heterozygous (HET) *MYH7* G256E hiPSC lines (G256E Cl.102, G256E Cl.141, G256E Cl.157) and two isogenic HET *MYH7* H251N hiPSC lines (H251N Cl.3, H251N Cl.85) were differentiated to hiPSC-derived CMs. All clones were harvested on day 30 post differentiation and three independent multiplexed scRNAseq runs were performed for each mutation using 10X Cell Plex system (Fig.1A). Differentiations yielded pure CM populations with high expression of CM markers *TNNT2* and *MYH7* and absent expression of endothelial (*ACE*) and fibroblast (*DCN*) markers for both mutations (Fig.S1A&B). Unbiased UMAP clustering of H251N CMs demonstrates the presence of 12 clusters with distinct separation between WT and HET H251N CMs (Fig. 1B-I). WT cells were primarily contributing (>70% normalized relative WT contribution) to clusters 1,3,4,5 while H251N mutant cells contributed mainly to clusters 2,6,7,8,9 & 10 (Fig.1B-II, III). Total *MYH7* expression was comparable in both WT and HET CMs (Fig.1B-IV). In contrast, unbiased UMAP clustering of G256E CMs lead to 12 clusters with no distinct separation of WT and HET mutant CMs with all 12 clusters showing comparable contribution (<60% relative normalized contribution) of WT and mutant cells except for one minor cluster (Fig.1C-I to III). Total *MYH7* expression was also comparable between WT and HET cells (Fig.1C-IV). DEG analysis revealed 165 DEGs for H251N HET and only 28 DEGs for G256E HET (Table-S1&2) vs WT hiPSC-CMs (log fold change > 0.5 & adjusted p-value < 0.05). H251N HET DEGs encompassed classic cardiac hypertrophy markers such as *NPPB* and *ANKRD1* as well as myocyte structural genes such *ACTN1* and genes involved in calcium handling such as *RYR2*. In addition to the hypertrophy related changes, GO-term cluster enrichment analysis also revealed changes in extracellular matrix remodeling in H251N CMs (Fig.S2A-C). In contrast, G256E DEGs showed mild upregulation of *NPPB* and *ANKRD1* as the only established cardiac hypertrophy markers. While a few skeletal muscle related genes were also regulated such as *TNNT1*, no overall expression pattern could be observed. GO-term analysis revealed no statistically significant enriched gene cluster (Fig.S3A&B).

**Fig. 1:**
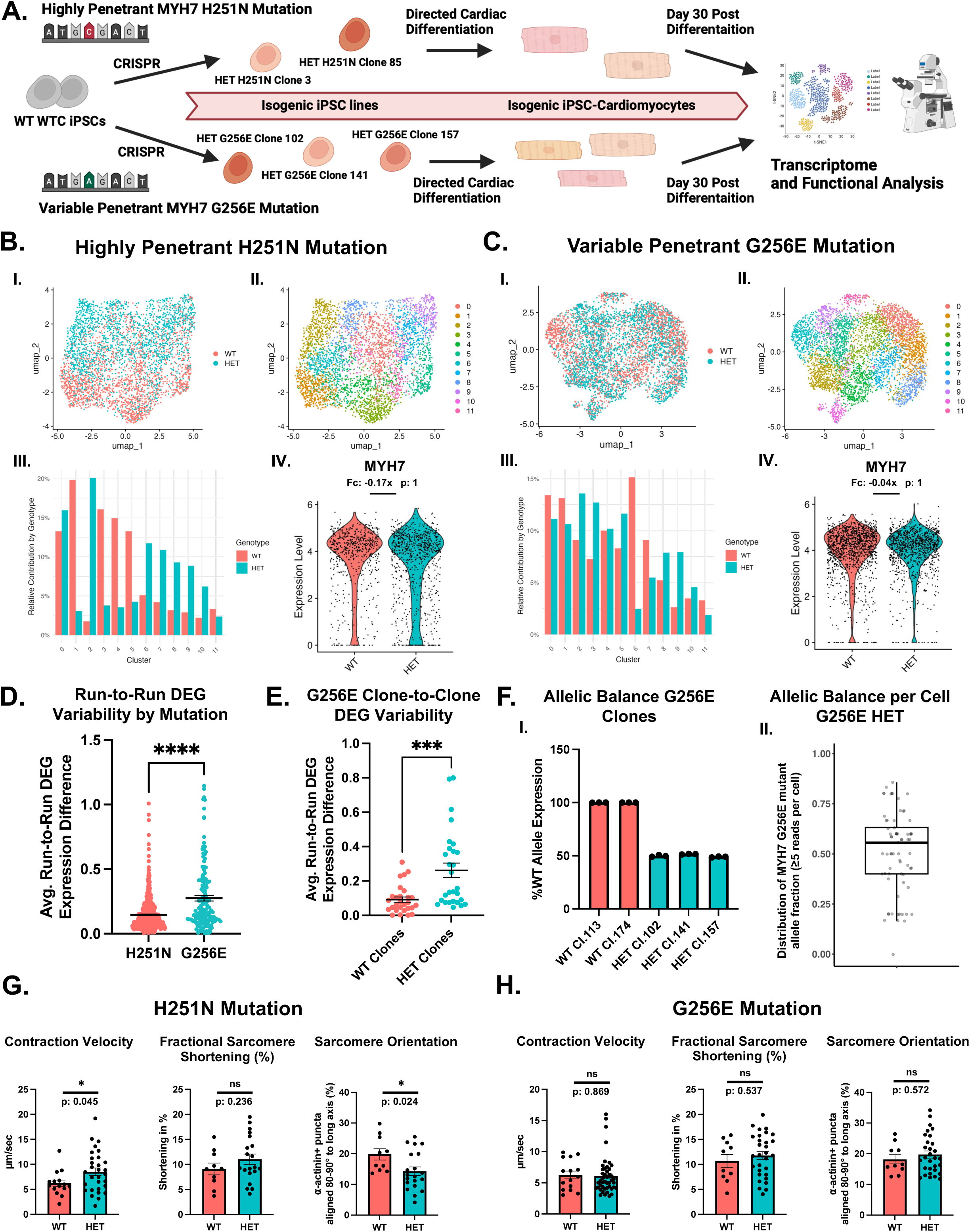
Distinct Phenotype Penetrance & Variability of Different *MYH7* Mutations is Reflected in hiPSC-CM Model. (A) Schematic of experimental set-up (B) I. UMAP representation of *MYH7* H251N CMs (*MYH7* WT/H251N) and the isogenic counterpart (*MYH7* WT/WT) clustered by the presence of H251N mutation. II. Unbiased clustering of H251N and WT hiPSC-CMs. III. Relative contribution to each cluster by WT and H251N hiPSC-CMs. IV. Comparison of *MYH7* gene expression in WT and H251N hiPSC-CMs. Fc = log2 fold change. (C) I. UMAP representation of *MYH7* G256E CMs (*MYH7* WT/G256E) and the isogenic counterpart (*MYH7* WT/WT) clustered by the presence of G256E mutation. II. Unbiased clustering of G256E and WT hiPSC-CMs. III. Relative contribution to each cluster by WT and G256E hiPSC-CMs. IV. Comparison of *MYH7* gene expression in WT and G256E hiPSC-CMs. Fc = log2 fold change. (D) Quantification of run-to-run expression variability of WT vs MUT DEGs between three independent scRNAseq runs. DEGs were defined as log fold change >0.5 & adjusted p-value < 0.05. Each dot represents the average run-to-run expression difference of one DEG for a specific clone. N(H251N) = 495 (3 clones, 165 DEGs); n(G256E) = 140 (5 clones, 28 DEGs). (E) Quantification of clone-to-clone expression variability of DEGs between three isogenic G256E HET CM clones and two isogenic WT CM clones. DEGs were defined as log fold change >0.5 & adjusted p-value < 0.05. Each dot represents the average clone-to-clone expression difference of one DEG. N = 28. (F) I. Quantification of WT allele expression via ddPCR for two isogenic WT CM clones and three isogenic G256E CM clones. Three biological replicates were analyzed per clone. II. Single cell mutant *MYH7* G256E vs. WT mRNA expression in two independent Kinnex scRNAseq runs of D30 HET Cl.141 CMs. (n=23, run1; n=46 run2). (G) Quantification of contraction velocity (CV), fractional sarcomere shortening (FSS) and sarcomere orientation (SO) of H251N HET (CV n=30, FSS&SO n=20, 2 isogenic clones) and WT CMs (CV n=15, FSS&SO n=10, 3 isogenic clones). (H) Quantification of CV, FSS and SO of G256E HET (CV n=45, FSS&SO n=30, 3 isogenic clones) and WT CMs (CV n=15, FSS&SO n=10, 3 isogenic clones). Data are presented as mean ± SEM. Statistical significance was determined by unpaired t test (E) and two-sided Wilcoxon rank-sum test with Bonferroni correction as implemented in Seurat (B,C). Functional parameters and Run-to-Run Variability were analyzed using linear mixed-effects models (lmerTest) with genotype as a fixed effect and clone as a random effect (Satterthwaite’s approximation) (D,G,H). *P < 0.05, **P < 0.01, ****P < 0.0001.

Importantly, H251N HET lines showed a significantly lower run-to-run variability in their DEGs when compared to the variability of DEGs in G256E runs (Fig.1D). Furthermore, in G256E CMs, HET clones showed a significantly higher clone-to-clone expression variability in their DEGs when compared to then their WT counterparts (Fig.1E). These observations were reflected in the unbiased clustering of CMs split by individual runs, where H251N CMs showed highly consistent clustering of each clone across all three runs whereas G256E CM clustering differed for each clone from run to run (Fig.S4A-C).

To investigate whether variable expression of the mutant *MYH7* allele in HET G256E lines represents an underlying cause of the variable DEGs in HET G256E CMs, we performed digital droplet PCR (ddPCR) to quantify the ratio of mutant and WT *MYH7* allele expression in hiPSC-CMs. While the two WT clones showed 100% WT allele expression across all three biological replicates, HET clones consistently showed a 1:1 expression ratio of mutant and WT alleles in all three isogenic lines across three independent biological replicates (Fig.1F-I & Fig.S5). To further assess allelic balance at single-cell resolution, we performed Kinnex long-read single-cell sequencing on G256E HET CMs and quantified the ratio of mutant versus WT reads per cell (Fig. S6A). While the bulk-level data recapitulated the ∼1:1 expression ratio, single-cell analysis revealed substantial heterogeneity, with individual cells ranging from 0% to 85.7% mutant *MYH7* allele expression (Fig. 1F-II & Fig. S6B). Finally, we examined whether cell-to-cell variability in mutant allele expression was associated with expression of previously identified mutation-associated markers. In the current dataset, this analysis did not provide sufficient evidence to establish a robust association between mutant mRNA fraction and mutation-associated marker expression (r = 0.08) (Fig. S6C,D). These findings suggest that, within the limits of the current sequencing depth, the relationship between allelic imbalance and transcriptional heterogeneity remains unresolved.

Functionally, sparsely plated H251N HET CMs exhibited a significantly higher contraction velocity compared to WT CMs (8.5 ± 0.7 μm/sec vs. 6.2 ± 0.6 μm/sec, p = 0.04), along with a significantly lower fraction of sarcomeres aligned with the long axis of the cell (14.3 ± 1.4% vs. 19.8 ± 1.8%, p = 0.02). In addition, we observed a trend toward increased fractional sarcomere shortening in H251N HET CMs (11.1 ± 1.0% vs. 9.1 ± 1.2%, p = 0.24). Together, these findings are consistent with a hypercontractile phenotype accompanied by structural disorganization (Fig. 1G & Fig. S7A&B). In contrast, G256E HET CMs did not exhibit a clear contractile or structural phenotype compared to WT controls (Fig. 1H & Fig. S7A&B).

### 3.2 Beta-Adrenergic Stimulation Reveals a Metabolic Stress Response Phenotype in Variably Penetrant *MYH7* G256E Mutation

We next investigated the impact of beta-adrenergic signaling-induced stress on HCM phenotype in *MYH7* G256E hiPSC-CMs by exposing them to 1 μM isoproterenol for five days (Fig.2A), following previously established protocols.^29,30^ Isoproterenol treatment resulted in a significant increase in beating rate and contraction velocity in both WT and HET G256E CMs, with no significant difference in contractile function observed between WT and HET G256E CMs post-treatment (Fig.2B). Two independent multiplexed scRNAseq runs of isoproterenol-treated CMs confirmed high expression of CM markers *MYH7* and *TNNT2*, while non-myocyte markers (*DCN*, *ACE*) were absent (Fig.S8A). Unbiased UMAP clustering of both WT and HET CMs from both runs identified 10 clusters (Fig.2C-I,II). Unlike in the non-isoproterenol-treated condition, isoproterenol treatment led to distinct contribution of WT into cluster 0 (83.0% normalized relative WT contribution) and HET into cluster 1 (88.6% normalized relative contribution). Clusters 2-9 displayed comparable contributions from each genotype (Fig. 2C-III). *MYH7* gene expression remained high in both WT and HET CMs, though a slight downregulation (log fold change: -0.16x; p = 1.6E-3) was observed in HET CMs (Fig.2C-IV). Analysis of DEGs between G256E HET vs WT CMs after isoproterenol treatment identified 49 DEGs. While 14 of the 49 DEGs were also significantly different between G256E HET vs WT at baseline without isoproterenol treatment, 35 DEGs showed significant difference only after isoproterenol treatment (Fig.2D & Table S3). GO-term analysis of the 49 identified DEGs revealed an enrichment of pathways associated with “protein kinase activity” “protein phosphorylation” and “ATP-binding” (Fig.S9A-C).

**Fig. 2:**
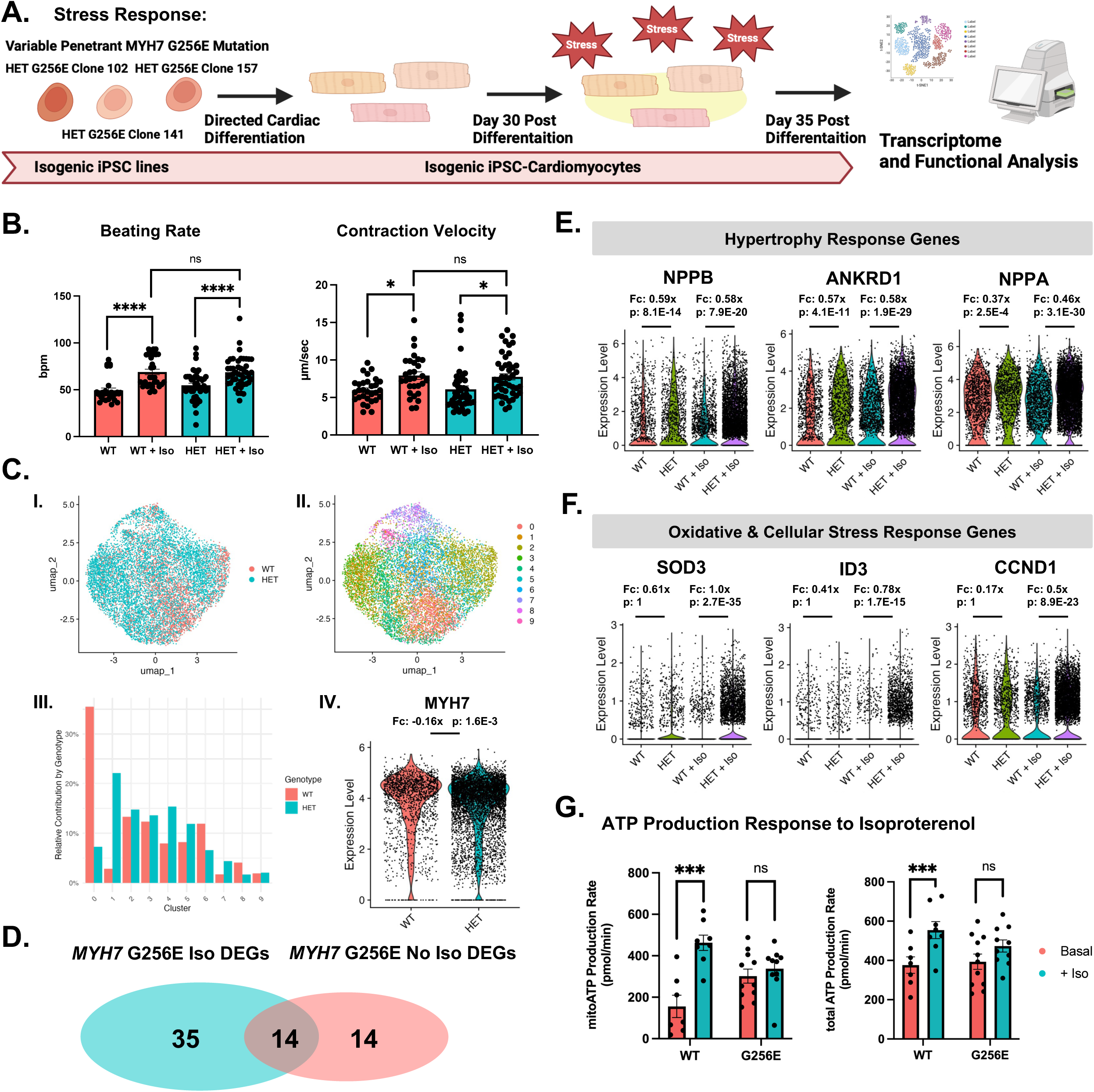
Beta-Adrenergic Stimulation Reveals a Metabolic Stress Response Phenotype in Variably Penetrant *MYH7* G256E Mutation. (A) Schematic of experimental set-up. (B) Quantification of beating rate and contraction velocity of WT (Basal&Iso: n=30, 2 isogenic clones) and *MYH7* G256E HET (Basal&Iso: n=45, 3 isogenic clones) CMs with and without isoproterenol (Iso) treatment. (C) I. UMAP representation of isoproterenol treated G256E CMs and the isogenic WT counterpart clustered by the presence of the G256E mutation. II. Unbiased clustering of G256E and WT hiPSC-CMs. III. Relative contribution to each cluster of WT and G256E hiPSC-CMs. IV. Comparison of *MYH7* gene expression in WT and G256E hiPSC-CMs. Fc = log2 fold change. (D) Venn diagram comparing WT vs. G256E DEGs of unstressed (No iso treatment) and stressed (Iso treatment) CMs. (E) Gene expression of established hypertrophy response genes in WT and G256E CMs with and without isoproterenol treatment (WT n=1332, HET n=1106, WT+Iso n=1604, HET+Iso n=5958). (F) Gene expression of oxidative and cellular stress response genes in WT and G256E CMs with and without isoproterenol treatment. Fc = log2 fold change. (G) Quantification of mitochondrial and total ATP production rate in WT vs. G256E CMs with and without isoproterenol treatment, measured by ATP Rate Assay (WT: basal n=7, iso n=8, 2 isogenic clones; HET: basal n=11, iso n=10, 3 isogenic clones). Data are presented as mean ± SEM. Statistical significance was determined by a two-sided Wilcoxon rank-sum test with Bonferroni correction as implemented in Seurat (C,E,F). Functional parameters were analyzed using linear mixed-effects models (lmerTest) with genotype as a fixed effect and clone as a random effect (Satterthwaite’s approximation) (B,G). *P < 0.05, **P < 0.01, ****P < 0.0001.

Hypertrophy markers *NPPB*, *ANKRD1* and *NPPA* exhibited comparable fold-change upregulation in HET vs WT CMs with beta-adrenergic stimulation compared to the baseline condition (Fig.2E). Interestingly, the differential expression of oxidative and protein folding stress response genes such as superoxide dismutase 3 (*SOD3*)^31^, inhibitor of DNA binding 3 (*ID3*)^32^, and Cyclin D1 (*CCND1*)^33^, between HET and WT CMs were increased following isoproterenol treatment but not at baseline (Fig.2F). Run-to-run variability in DEGs was significantly reduced in isoproterenol-treated CMs compared with baseline conditions, and clone-to-clone variability was not increased in HET CMs compared to WT CMs, as previously observed in the baseline condition (Fig. S8B–D). We subsequently examined the level of cellular respiration in isoproterenol-treated G256E HET CMs using the Seahorse assay and confirmed that G256E HET CMs exhibited elevated mitochondrial respiration and ATP production at baseline.^18^ With isoproterenol treatment, WT CMs appropriately upregulated their mitochondrial respiration and ATP production, while G256E HET CMs failed to do so (Fig.2G). Significant variation in glycolysis-dependent ATP production was also noted (Fig.S10).

### 3.3 Increase in Mutant Gene Dosage Results in Consistent Change in Transcriptional Signature

Based on previously reported single-cell analyses demonstrating substantial allelic imbalance^9,10,11^ as well as our own long-read–based quantification of allele-specific expression ratios (Fig. 1F-II), we next investigated the impact of increased mutant gene dosage on phenotype penetrance in G256E CMs. We therefore engineered two isogenic cell lines harboring a homozygous (HOM) *MYH7* G256E mutation (G256E Cl.15 & Cl.17), modeling the upper extreme of the allelic imbalance spectrum (Fig. 3A). Successful gene editing was confirmed via Sanger sequencing (Fig.3B). The HOM *MYH7* G256E hiPSCs were then differentiated into day 30 CMs, and two independent multiplexed scRNAseq experiments were conducted to assess overall transcriptomic changes. We confirmed the high efficiency of CM differentiation (Fig.S11A) and identified eight distinct WT and HOM CM clusters following UMAP dimensionality reduction (Fig. 3C-I,II). In contrast to G256E HET CMs that showed no difference in clustering from WT CMs (Fig. 1C), HOM CMs and WT CMs exhibited significantly different clustering patterns, with WT CMs found predominantly in clusters 1, 5, 6, and 7 (>79% relative normalized WT contribution) and HOM CMs predominantly in clusters 2, 3, and 4 (>80% relative normalized HOM contribution) (Fig. 3C-III). Both groups expressed high levels of *MYH7*, albeit with slightly lower expression in HOM CMs (log fold change: -0.39x; p = 2.9E-13) (Fig.3C-IV). Analysis of HOM vs WT gene expression identified 260 DEGs, with 23 of these DEGs overlapping with DEGs from the HET vs WT analysis (Fig. 3D, Fig.S12A & B & Table S4). Gene ontology analysis revealed an enrichment in cytoskeletal functions such as “actin binding” and “actin filament organization” as well as ECM remodeling (“collagen binding” & “cell-matrix adhesion” and calcium cycling (“calcium ion binding” & “ion channel binding”) (Fig.S12C). The run-to-run and clone-to-clone variability in DEG expression levels was significantly lower in HOM CMs lines compared to HET CMs (Fig.3E,F).

**Fig. 3:**
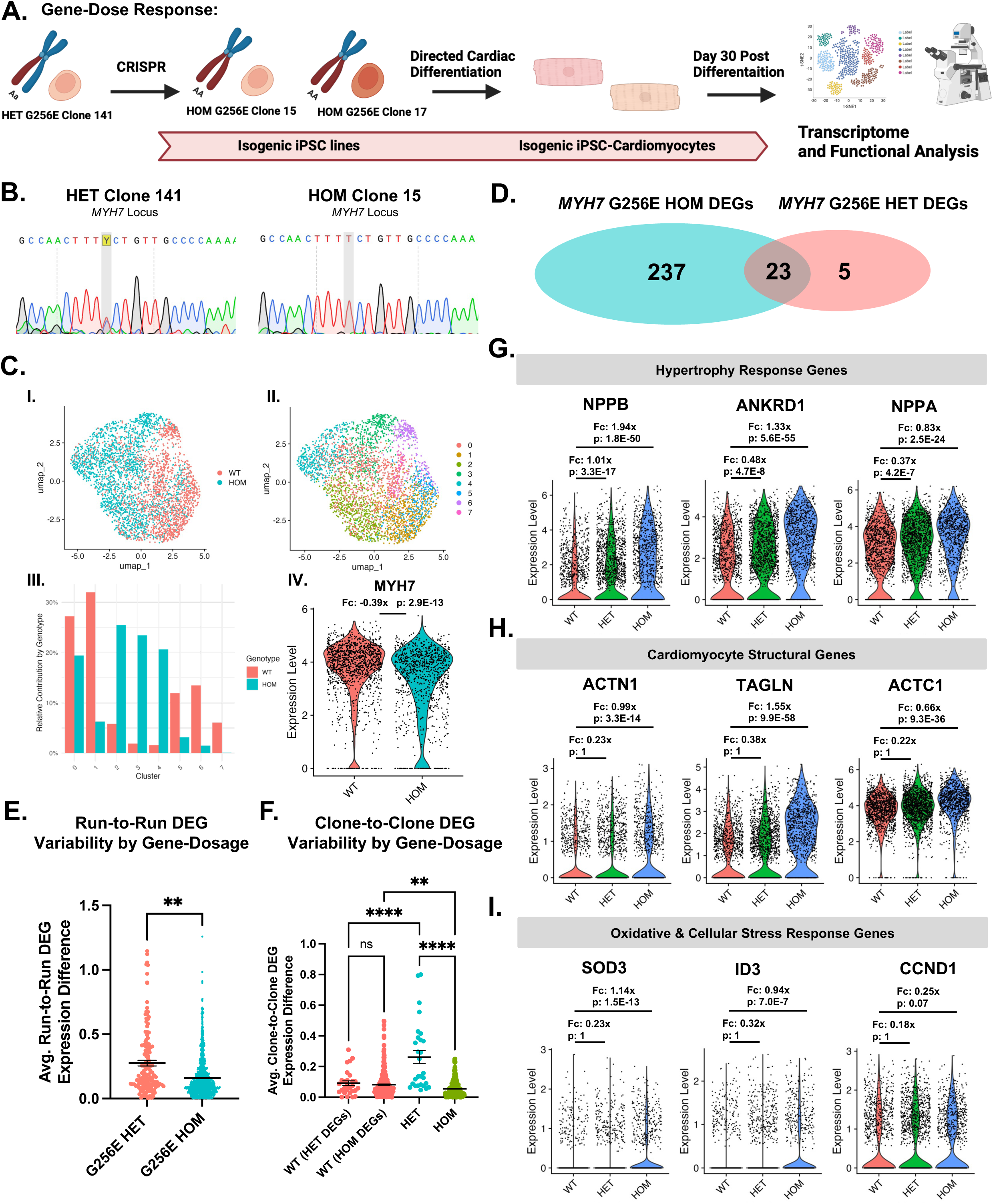
Increase in Mutant Gene Dosage Results in Consistent Change in Transcriptional Signature. (A) Schematic of experimental set-up. (B) Sanger Sequencing of *MYH7* locus at G256E mutation site of HET clone 141 and HOM clone 15. (C) I. UMAP representation of HOM *MYH7* G256E mutant cells (*MYH7* G256E/G256E) and the isogenic WT counterpart (*MYH7* WT/WT) clustered by the presence of G256E mutation. II. Unbiased clustering of G256E and WT hiPSC-CMs. III. Relative contribution to each cluster by WT and G256E HOM hiPSC-CMs. IV. Comparison of *MYH7* gene expression in WT and G256E HOM hiPSC-CMs. Fc = log2 fold change. (D) Venn diagram comparing WT vs G256E HOM DEGs with WT vs. G256E HET DEGs. (E) Quantification of run-to-run expression variability of DEGs between independent scRNAseq runs. DEGs were defined as log fold change >0.5 & adjusted p-value < 0.05. Each dot represents the average run-to-run expression difference of one DEG for a specific clone. N(G256E HET) = 140 (5 clones, 28 DEGs); n(G256E HOM) = 1040 (4 clones, 260 DEGs). (F) Quantification of clone-to-clone expression variability of DEGs between two isogenic G256E HOM CM clones, three isogenic G256E HET CM clones and two isogenic WT CM clones. DEGs were defined as log fold change >0.5 & adjusted p-value < 0.05. Each dot represents the average clone-to-clone expression difference of one DEG. N(HET) = 28, n(HOM) = 260. (G-I) Gene expression of established hypertrophy response genes (G), myocyte structural genes (H) and oxidative & cellular stress response genes (I) in baseline WT and G256E HET & HOM CMs (WT n=1304, HET n=1620, HOM n=980). Fc = log2 fold change. Data are presented as mean ± SEM. Statistical significance was determined by linear mixed-effects models (lmerTest) with genotype as a fixed effect and clone as a random effect (E), ordinary one-way ANOVA with post-hoc Tukey’s multiple comparisons test (F) and a two-sided Wilcoxon rank-sum test with Bonferroni correction as implemented in Seurat (C,G,H,I). \**P* < 0.05, \*\**P* < 0.01, \*\*\*\**P* < 0.0001.

Hypertrophy markers *NPPB*, *ANKRD1*, and *NPPA* exhibited a strong gene dose-dependent expression pattern, with the lowest expression in WT CMs and highest in HOM CMs (Fig.3G). Additionally, sarcomeric protein genes such as Alpha-Actinin 1 (*ACTN1*), Transgelin (*TAGLN*) and Cardiac Alpha-Actin 1 (*ACTC1*), which were only mildly upregulated in HET CMs showed strong upregulation in HOM CMs (Fig.3H). Interestingly, genes that were upregulated with isoproterenol treatment (*SOD3*, *ID3* and *CCND1)* showed only mild (*SOD3*, *ID3*) to no upregulation (*CCND1*) in HOM CMs (Fig.3I).

### 3.4 Proteomic Profiling Confirms Gene Dose-Dependent Transcriptional Changes at the Protein Level

Given the significant transcriptomic alterations observed with increased gene dosage, we proceeded to examine proteomic changes in *MYH7* G256E HET and HOM CMs. Utilizing TMTpro multiplexing, we analyzed two WT, two HET, and two HOM cell lines, each differentiated into CMs and evaluated by mass spectrometry in biological triplicates (Fig. 4A). Our analysis yielded a total of 345,425 peptide-spectrum matches (PSMs), corresponding to 94,882 distinct peptides from 8,229 proteins (Fig. 4B). Quantification of the protein abundances confirmed similar levels of MYH7 across all lines (Fig. S13A).

**Fig. 4:**
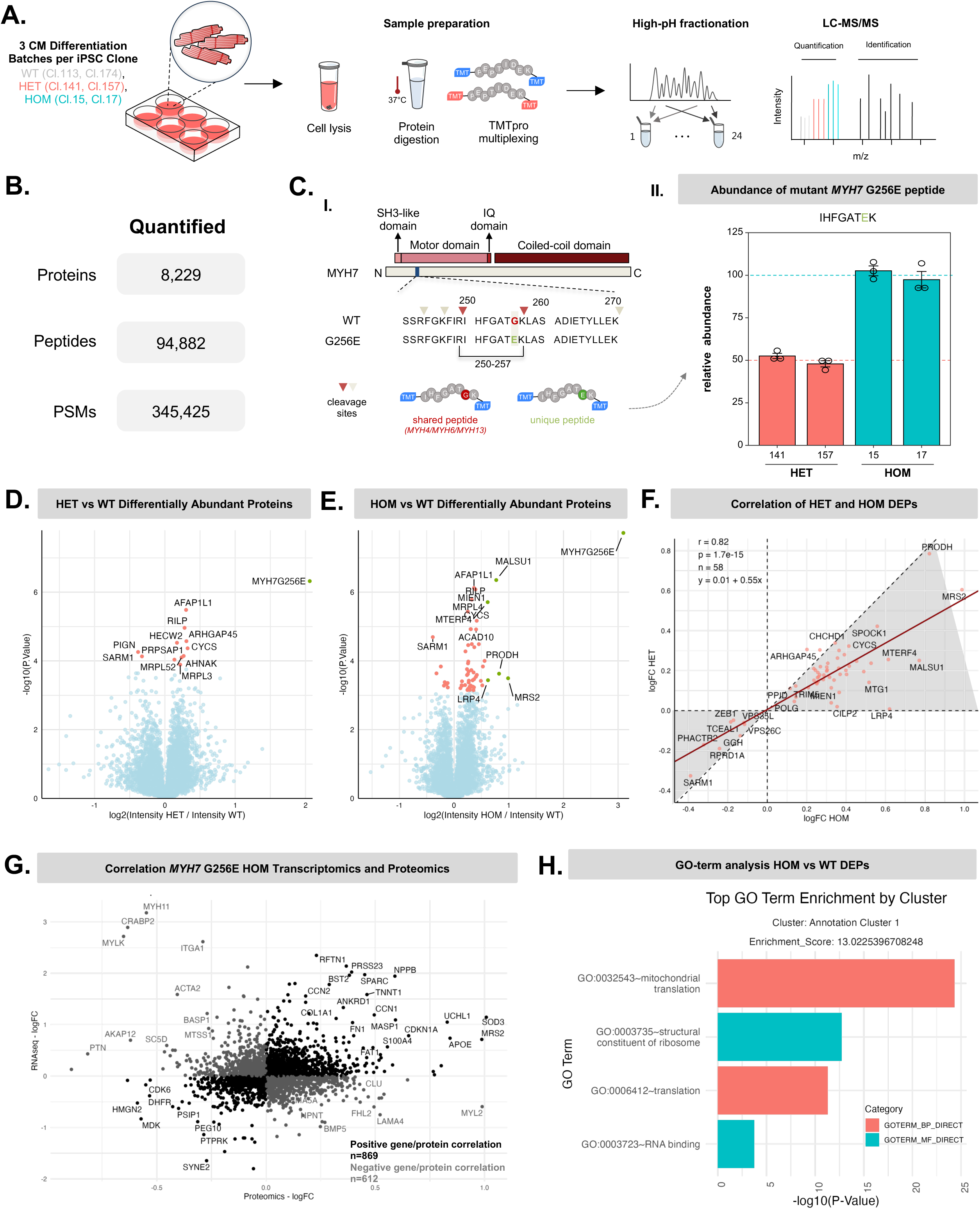
Proteomic Profiling Confirms Gene Dose Dependent Transcriptional Changes at the Protein Level. (A) Schematic of experimental set-up. Three biological replicates were quantified per clone (WT Cl.113 & Cl.174, HET Cl.141 & Cl.157, HOM Cl.15 & Cl.17) (B) Number of features quantified in the TMT experiment. (C) Quantification of MYH7 proteoforms in isogenic G256E HET and HOM CM clones. I. Schematic of the myosin heavy chain 7 sequence. Wild-type and G256E MYH7 proteoforms are differentiated based on the tryptic peptide spanning amino acids 250-257. II. Quantification of the mutant MYH7 G256E peptide in HET and HOM CMs. Data are presented as mean ± SE. (D+E) Volcano plot of differently abundant proteins in G256E HET vs. WT CMs (D) and G256E HOM vs. WT CMs (E). Salmon color indicating statistical significance defined as p-value < 0.05 and false discovery rate (FDR) < 0.1. Green color indicating both statistical significance and effect size of log2-fold change > 0.58 (F) Scatterplot comparing HET vs WT and HOM vs WT fold change regulation of previously identified HOM vs WT DEPs (n=58). Grey area indicates gene-dose dependent regulation of DEPs, red line represents linear regression fit. (G) Correlation of G256E HOM vs WT fold changes in proteomic and transcriptomic dataset. Black color indicating positive gene/protein expression/abundance correlation & grey color indicating negative gene/protein expression/abundance correlation (H) Top enriched GO-terms for identified G256E HOM vs WT DEPs.

To evaluate the allelic abundance of WT and mutant MYH7 in the CMs, we quantified the peptide IHFGATEK, which is specifically derived from the G256E mutation, in both HET and HOM CMs (Fig.4C-I). In HET CMs, the abundance of the mutant peptide was 50% of that observed in HOM CMs, after normalization to the total MYH7 content. This indicates that mutant peptide levels in HET CMs correlate directly with the number of mutant alleles, reflecting an expected 50:50 abundance ratio of mutant to wild-type MYH7 isoforms (Fig. 4C-II, Fig. S13A). This result aligns with our ddPCR data for G256E HET CMs mRNA expression.

Differential protein abundance analysis identified 11 proteins with statistically significant abundance changes between WT and G256E HET CMs, none of which exceeded an absolute log2 fold-change threshold of 0.58, corresponding to approximately 1.5-fold change. In G256E HOM versus WT CMs, 58 proteins showed statistically significant abundance changes, of which five also exceeded the 1.5-fold abundance threshold: MALSU1, MTERF4, PRODH, MRS2, and LRP4 (Fig. 4D,E; Fig. S13B; Tables S5,S6; BH-adjusted p-value < 0.1). To assess whether protein abundance changes followed a gene-dose-dependent pattern, we compared log2 fold-change values between G256E HOM and G256E HET CMs for the 58 proteins that were differentially abundant in G256E HOM versus WT CMs. HET and HOM abundance changes were strongly correlated by Spearman correlation analysis (ρ = 0.82, p = 1.7 × 10⁻¹⁵, n = 58). Linear regression yielded a slope of 0.55, consistent with attenuated abundance changes in HET compared with HOM CMs. At the individual protein level, 56 of 58 proteins showed a gene-dose-dependent abundance pattern, defined as regulation in the same direction in HET and HOM with the HET log2 fold-change closer to zero than the corresponding HOM log2 fold-change (Fig. 4F; Fig. S13B).

Comparison of RNA-level and protein-abundance changes revealed modest transcriptome–proteome concordance. Specifically, among all quantified proteins with corresponding transcript measurements, 52.7% in HET CMs and 58.7% in HOM CMs changed in the same direction as their corresponding transcripts (Fig. 4G and Fig. S13C). Gene ontology enrichment analysis of the most differentially regulated proteins revealed enrichment of terms related to mitochondrial translation, structural constituents of the ribosome, and translation (Fig. 4H).

### 3.5 Homozygosity Reveals a Hypercontractile Phenotype in Variably Penetrant HCM Mutation

Given the gene dose–dependent increase in transcriptomic and proteomic phenotypes, we next assessed whether contractile and structural features were more consistently altered in G256E HOM compared to G256E HET CMs. Motion analysis of sparsely plated cells revealed a significant increase in contraction velocity (5.9 ± 0.6 μm/s in WT vs. 8.4 ± 0.7 μm/s in HOM, p = 0.02), accompanied by higher fractional sarcomere shortening (7.9 ± 1.0% in WT vs. 15.7 ± 0.9% in HOM, p < 0.01). In parallel, HOM CMs exhibited reduced alignment of sarcomeres with the long axis of the cell (17.4 ± 1.5% in WT vs. 13.0 ± 1.0% in HOM, p = 0.01) (Fig. 5A & B). The relaxed Z-to-Z distance remained unchanged between WT and HOM cells. Collectively, these data indicate that increased mutant gene dosage in *MYH7* G256E CMs results in a hypercontractile and structurally disorganized phenotype that is not observed in G256E HET cells.

**Fig. 5:**
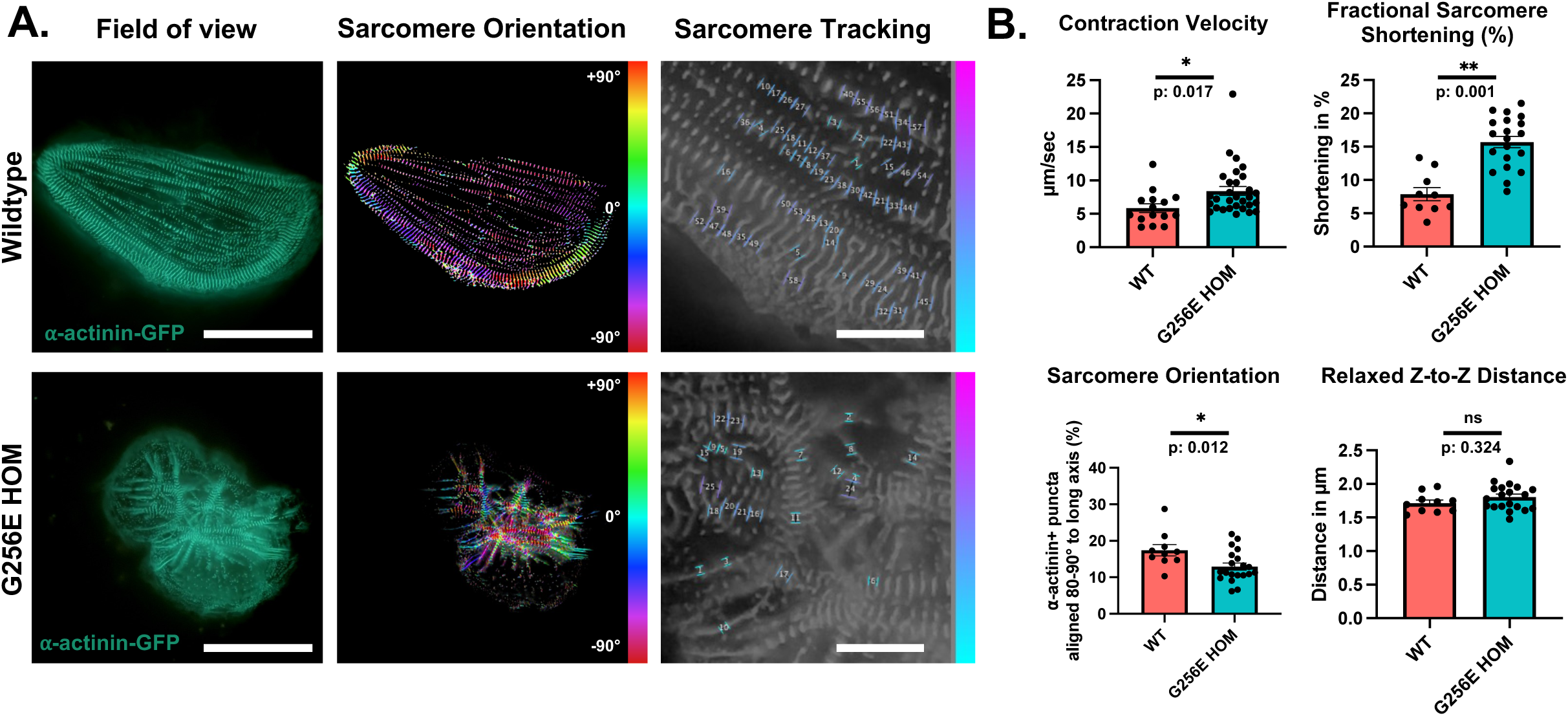
Homozygosity Reveals a Hypercontractile Phenotype in Variably Penetrant HCM Mutation. (A) Representative images of WT and G256E HOM cardiomyocytes showing, from left to right, α-actinin–GFP fluorescence, sarcomere orientation (color indicates local sarcomere orientation relative to the horizontal axis [cyan = horizontal, red = ±90°, vertical]), and sarcomere tracking (color-coded by Z-to-Z distance from longest [pink] to shortest [blue]). Scale bars: 50 μm (field of view, sarcomere orientation), 10 μm (sarcomere tracking). (B) Quantification of contraction velocity (CV), fractional sarcomere shortening (FSS), sarcomere orientation (SO), and relaxed Z-to-Z distance (RZ) in G256E HOM (CV n = 30; FSS, SO, RZ n = 20; 2 isogenic clones) and WT cardiomyocytes (CV n = 15; FSS, SO, RZ n = 10; 3 isogenic clones). Data are presented as mean ± SEM; functional parameters were analyzed using linear mixed-effects models (lmerTest) with genotype as a fixed effect and clone as a random effect (Satterthwaite’s approximation) (B). *P < 0.05, **P < 0.01, ****P < 0.0001.

### 3.6 Identification of Common vs Mutation-specific HCM Markers

To address whether the transcriptome phenotype shift of G256E mutant hiPSC-CMs in response to isoproterenol treatment is similar to that with increasing the mutant gene dosage, we combined and compared scRNAseq data from all experimental conditions (*MYH7* G256E HET and HOM CMs, isoproterenol-treated *MYH7* G256E HET CMs, and *MYH7* H251N HET CMs) (Fig.6A). DEG integration revealed six genes that were consistently upregulated across all conditions: *NPPB*, *PDLIM3*, *APOE*, *ANKRD1*, *RAI14* and *SPARC* (Fig.6B). DEG integration also revealed a high number of DEGs specific to the G256E HOM condition (157 DEGs) and the H251N HET condition (77 DEGs), as well as 70 DEGs shared between just these two experimental conditions (Fig.6A). Another six genes were regulated consistently in all G256E experimental conditions, but not in H251N CMs (upregulated: *CDKN1A*, *HOPX*, *TNNT1*; downregulated: *PEG10*, *SYNE2*, *LDHA*) (Fig.S14A). GO-term analysis of mutation-specific DEGs revealed that H251N specific DEGs were mostly related to cell adhesion remodeling (“cell adhesion”, “collagen binding”, “integrin binding”) while G256E HOM specific DEGs were related to changes in the contractile apparatus (“actin filament binding”, “actin binding”). The shared DEGs between these two conditions were mostly involved in ECM remodeling (“extracellular matrix structural constituent”) (Fig.6C-E & Table S7).

**Fig. 6:**
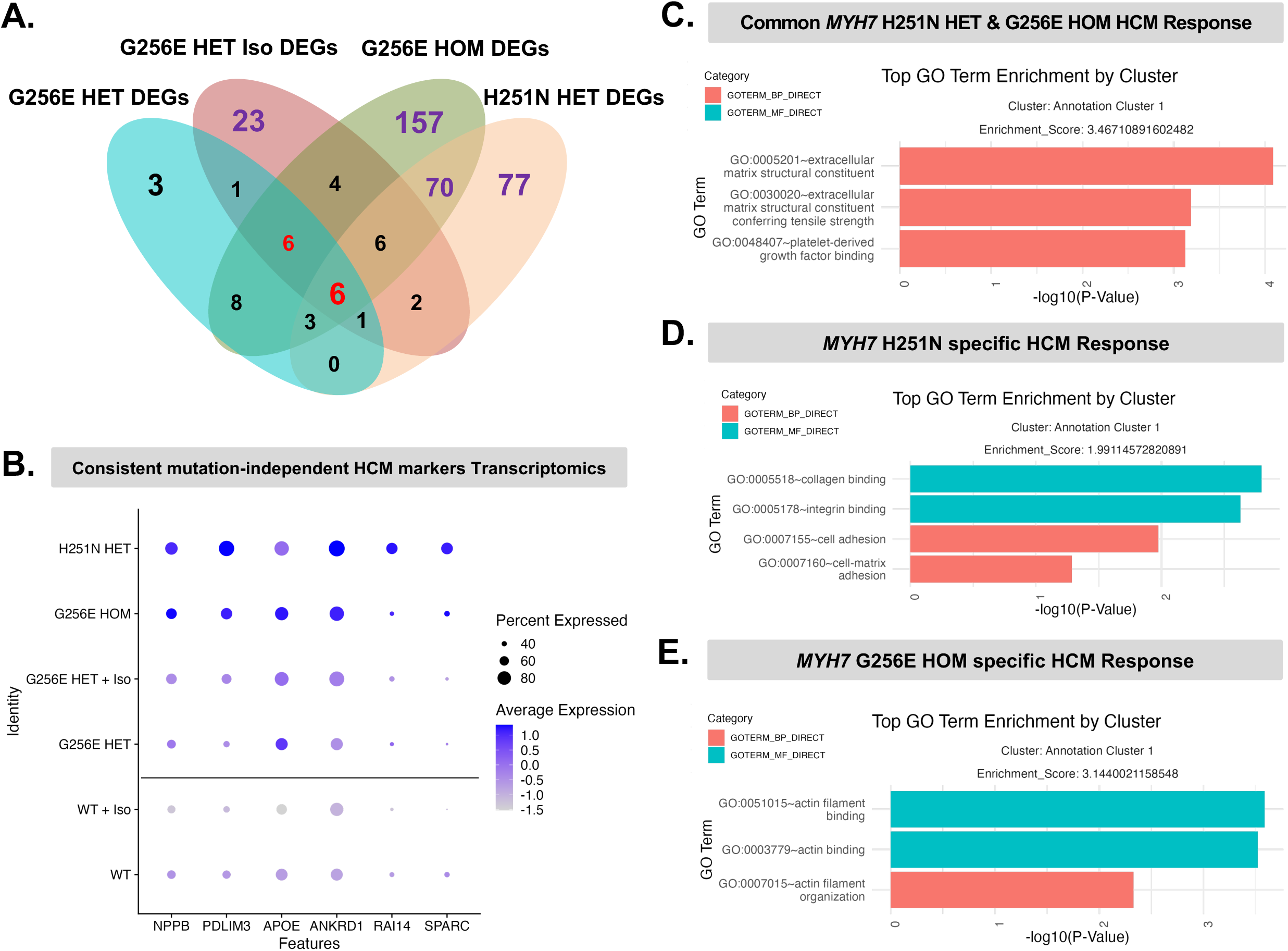
Identification of Common vs Mutation-specific HCM Markers. (A) Venn diagram of DEGs across all four experimental conditions. (B) Dotplot expression of identified markers consistently upregulated in all G256E & H251N transcriptomic experiments. (C-E) Top GO-term enrichment clusters for identified DEGs specific to only G256E HOM and H251N HET CMs (C), only H251N HET CMs (D) and only G256E HOM CMs (E).

We subsequently assessed the protein abundance of previously identified markers that were consistently altered at the transcriptional level across all conditions. Among these, NPPB and ANKRD1 exhibited gene dose–dependent abundance trends in the proteomics dataset, which were further evaluated by independent Western blot analysis (Fig. S15A&B).

## 4. Discussion

Despite the identification of over 200 *MYH7* mutations, the phenotypic presentation of HCM remains highly variable, not only among different mutations but also among patients carrying the same HCM-associated mutation.^6,8^ This variability complicates the prediction and management of the disease considerably. This study aimed to elucidate the factors contributing to this variability and drivers of phenotype penetrance in variably penetrant mutations. We contrasted the highly penetrant *MYH7* H251N mutation, known for its strong early-onset HCM phenotype both clinically and in cell culture assays^4,19^ with the *MYH7* G256E mutation that exhibits a range of clinical phenotypes^6^ and a mild phenotype in vitro^18^, to address factors that drive phenotype penetrance. Using genome-edited isogenic hiPSC-CMs, we assessed the effects of the gene mutations without the confounding influence of background genetic modifiers that would be present using patient-derived hiPSCs. We increased the rigor of our findings by performing studies using multiple clones for each mutation and genotype, as well as multiple independent scRNAseq transcriptomic experiments, supplemented by proteomics and functional studies. This comprehensive approach allowed us to identify specific factors driving the variability in phenotype penetrance, enhancing our understanding of the underlying mechanisms in HCM.

Our primary findings reveal that both beta-adrenergic stimulation and increasing gene dosage significantly enhance the penetrance of disease phenotypes in variably penetrant *MYH7* G256E hiPSC-CMs. However, these factors induce two distinct phenotype changes. Specifically, beta-adrenergic stimulation leads to increased oxidative phosphorylation and a failure of G256E HET CMs to enhance mitochondrial ATP production. On the other hand, increasing gene dosage resulted in a pronounced hypercontractile and structurally disorganized phenotype in HOM *MYH7* G256E CMs that was absent in HET CMs. In addition, HOM *MYH7* G256E CMs showed an elevated expression of myocyte structural genes on a transcriptomic level and an increase in proteins related to mitochondrial metabolism and protein translation. DEG analysis across various conditions further identified a set of HCM marker genes that were consistently regulated in all mutant lines along with genes that were specific to only one *MYH7* mutation.

Given the known phenotypic variability of the MYH7 G256E mutation, we analyzed multiple isogenic clones across several scRNAseq runs, recognizing the susceptibility of scRNAseq to increased false discovery rates.^34^ Integrating runs with the Harmony algorithm,^23^ we found a clear clustering distinction between highly penetrant H251N CMs and WT CMs, while *MYH7* G256E CMs did not separately cluster from WT CMs. These findings support the use of UMAP clustering as an additional approach to assess the consistency of transcriptomic phenotypes associated with variable penetrance, while also identifying cluster-specific transcriptional signatures across genotypes and conditions (Fig. S16). Although batch integration methodologies are well-established for cell embedding, DEG calculations remain susceptible to batch effects, with optimal correction strategies still under debate.^35^ To address batch confounding in DEG analysis, we down sampled each dataset to achieve comparable cell counts per clone and run, enhancing the reliability of our results.

Upon establishing isogenic CMs as a model for studying HCM phenotype penetrance, we assessed the response to beta-adrenergic stimulation in G256E CMs. Clinically, adrenergic stress is associated with left ventricular hypertrophy in HCM patients,^36^ while beta-blockers improve symptoms and survival.^14^ Our analysis revealed a distinct metabolic response in HCM CMs subjected to beta-adrenergic stimulation, marked by an increased expression of oxidative stress genes and an impaired capacity to upregulate mitochondrial ATP production. While mitochondrial dysfunction in HCM is well-documented,^37^ our findings indicate a metabolic bottleneck that could exacerbate HCM pathophysiology under beta-adrenergic stimulation, such as occurs during exercise. WT CMs have lower baseline ATP production and can increase ATP in response to beta-adrenergic stimulation, whereas HCM CMs have higher baseline ATP production but fail to upregulate it after beta-adrenergic stimulation. These findings underscore the critical role of mitochondrial reserve capacity in beta-adrenergic stimulation, revealing a novel aspect of metabolic vulnerability in HCM CMs.

We next investigated the impact of gene dosage on HCM phenotype penetrance by generating two HOM *MYH7* G256E cell lines. Our study revealed that transcriptomic and proteomic phenotype alterations in G256E CMs are highly responsive to increased gene dosage. Specifically, we observed that higher gene dosage increased the mild changes observed in HET CMs rather than eliciting a separate distinct phenotype, as seen during beta-adrenergic stimulation. Additionally, increasing gene dosage significantly reduced clone-to-clone and run-to-run variability, suggesting utility of HOM models for studying variable and low-penetrant disease mechanisms. The pronounced transcriptomic, proteomic, and functional changes in response to gene dosage, in combination with the observed allelic imbalance at the single-cell level, warrant further studies of the temporal dynamics of allelic balance and the translation of allelic imbalance into cellular phenotypes, for example using bi-allelic reporter systems. Furthermore, our results indicate that further research on modifying allelic balance through allele-specific silencing^38^ and preventing transcriptional bursts^11^ could represent promising strategies to influence HCM disease penetrance.

Intersecting our datasets identified a set of consistent mutation-associated transcriptomic candidate markers, including NPPB, PDLIM3, APOE, ANKRD1, RAI14, and SPARC. Cross-referencing these candidates with a human proteomic dataset derived from myocardial samples of patients with obstructive HCM showed that, with the exception of RAI14, these markers were also significantly increased at the protein level in human HCM myocardium.^39^ The strongest and most significant changes were observed for ANKRD1 and NPPB, with ANKRD1 showing a 2.56-fold increase in abundance (*p* = 7.90 × 10⁻⁶) and NPPB showing a 1.76-fold increase (*p* = 0.002), supporting the relevance of these markers in human HCM tissue. More modest but statistically significant increases were observed for PDLIM3 (1.26-fold, *p* = 0.041), APOE isoforms E3 and E4 (1.31-fold and 1.21-fold, *p* = 0.017 and *p* = 0.043, respectively), and SPARC (1.17-fold, *p* = 0.021).^39^ Further studies will be required to define the functional roles and cellular origins of these genes and their corresponding proteins in the context of HCM.

### 4.1 Study Limitations

While our study provides important insights into phenotype penetrance in HCM, it is not without limitations. First, although hiPSC-CMs represent a powerful and widely used model system, they do not fully recapitulate the complex in vivo environment of the human heart. Differentiation protocols and culture conditions can introduce variability and yield cells that remain structurally and metabolically immature compared to adult CMs. To partially address this limitation, we performed additional validation experiments using day 30 G256E HET hiPSC-CMs cultured in a previously defined maturation medium^40^, as well as day 60 G256E HET CMs. These analyses showed that transcriptional variability associated with the lowly penetrant G256E mutation remained abundant in these more mature culture conditions. Moreover, the more mature G256E HET CMs did not acquire the stronger transcriptional phenotypes observed in H251N or G256E HOM cells, supporting that the variability reflects, at least in part, the heterozygous G256E variant context rather than developmental immaturity alone (Fig. S17 & S18). Additionally, prior studies have shown that key functional features are established in the day 30 hiPSC-CMs analyzed here. This includes a MYH7/MYH6 expression ratio approaching that of adult CMs and β-adrenergic receptor expression and functional downstream signaling established by day 30 of differentiation. At this stage, signaling is predominantly mediated by β₂-adrenergic receptors, with a shift toward β₁-adrenergic receptor dominance as the cells mature.^30^ Furthermore, for metabolic analyses hiPSC-CMs were maintained in glucose-rich media, which does not fully reflect the fatty acid–dependent metabolism of adult CMs. While this may influence mitochondrial capacity and stress responses, our findings highlight a relative impairment in metabolic adaptability of mutant cells under β-adrenergic stimulation. Future studies incorporating fatty acid supplementation will be important to determine how these phenotypes translate to more physiologically relevant metabolic conditions. Finally, while our study integrates transcriptomic, proteomic, structural, and metabolic analyses, further expansion of functional phenotyping, particularly with respect to calcium handling and electrophysiology in more mature hiPSC-CMs, will be important, as these features may become more pronounced at later stages of maturation.^41,42^

### 4.2 Future Studies

We plan to expand the number of different *MYH7* mutations in our studies to gain a broader understanding of the factors influencing HCM phenotypic variability. Incorporating more advanced 3D cardiac tissue models and in vivo validation studies will also help to confirm the findings from in vitro hiPSC-CMs studies and further reveal the functional consequences of these mutations in the context of whole heart physiology. Investigating the role of additional genetic and non-genetic modifiers in HCM penetrance will also be important. This includes exploring the impact of other sarcomere protein mutations, epigenetic modifications, and environmental factors such as diet and lifestyle. In conclusion, our study establishes a new avenue in hiPSC-based disease modelling by utilizing a variable penetrant mutation to investigate driving factors of phenotype penetrance. Specifically, our study highlights the critical influence of gene dosage and beta-adrenergic stress on development of distinct HCM disease-associated phenotypes, providing novel insights into mechanisms that may shape variable disease expression in HCM.

## Supporting information

Manuscript_Supplement

## Funding

Boehringer Ingelheim Fonds MD Fellowship and AHA Predoctoral Fellowship (23PRE1018158) (to P.H.); National Research Foundation of Korea grant funded by the Ministry of Education (RS-2022-NR070873), and the Ministry of Science and ICT (RS-2024-00342554), and Korea Health Industry Development Institute (KHIDI), funded by the Ministry of Health & Welfare (RS-2022-KH129482, RS-2024-00437518) (to S.L.); grants from the Novo Nordisk Foundation (NNF20OC0059767 to A.L., NNF20SA0064340 - BRIDGE Translational Excellence Programme (bridge.ku.dk) to C.S.), a PhD research grant from the Danish Cardiovascular Academy (PhD2023011-DCA to J.S.A.), which is funded by the Novo Nordisk Foundation (NNF20SA0067242); European Research Council (ERC grants 788381 and 101141820), the German Research Foundation (TRR_152 and TRR_267), and the German Centre for Cardiovascular Research (DZHK 81Z0600601) (to A.M.); NIH K99HL153679 (to A.V.R); H.Z. has received support from National Institutes of Health grants 1K08HL16140501, R01HL174432, 1R01HL177581-01A1, and R03HL173146; the AHA Transformational Project Award 25TPA1480322; the Sarnoff Scholar Award; and the Stanford CVI Seed grant. American Heart Association - Established Investigator Award, Hoffmann/Schroepfer Foundation, NIH/NIGMS (RM1 GM131981), Additional Venture Foundation, Joan and Sanford I. Weill Scholar Fund, and the NSF RECODE grant (to S.M.W)

## Author Contribution Statement

P.H., S.L., R.J., J.S.A., V.N., C.L., C.S., H.D., A.V.R, F.S., J.J., A.K., D.L., A.P., and B.R. performed research and analyzed data. P.H., S.L., J.S.A, H.N, H.Z., J.W., D.B, A.M., A.L. and S.W. edited the paper; P.H., S.L., J.S.A, A.L. and S.W. designed the research and wrote the paper.

## Acknowledgments

We thank members of the Wu lab and members of the NIGM RM1 investigative group for their comments and critiques throughout the development of this manuscript. Mass spectrometry analyses were performed at the Proteomics Research Infrastructure at the University of Copenhagen, supported by the Novo Nordisk Foundation (NNF19SA0059305). Sarcomere imaging was performed using the Stanford Neuroscience Microscopy Service. Figure schematics were generated using BioRender.com.

## Conflict of Interest

None

## SUPPLEMENT

**Supplement Figure 1.**
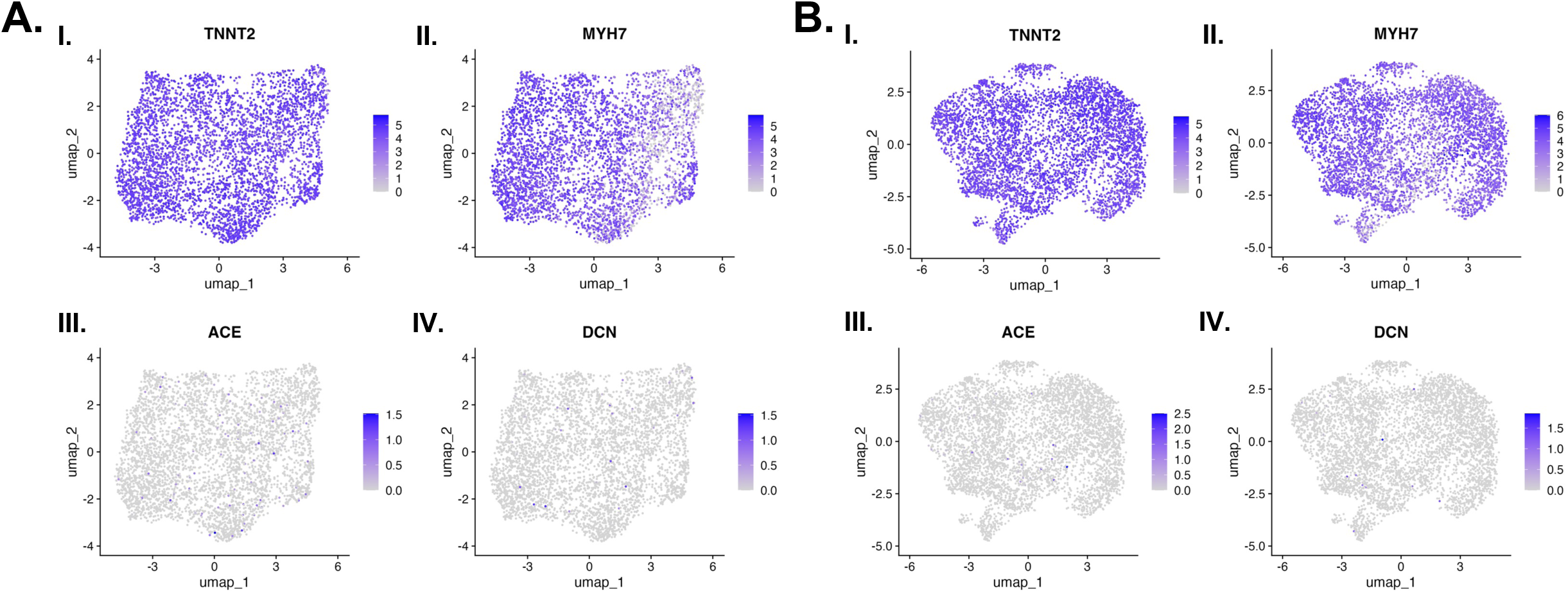

**Supplement Figure 2.**
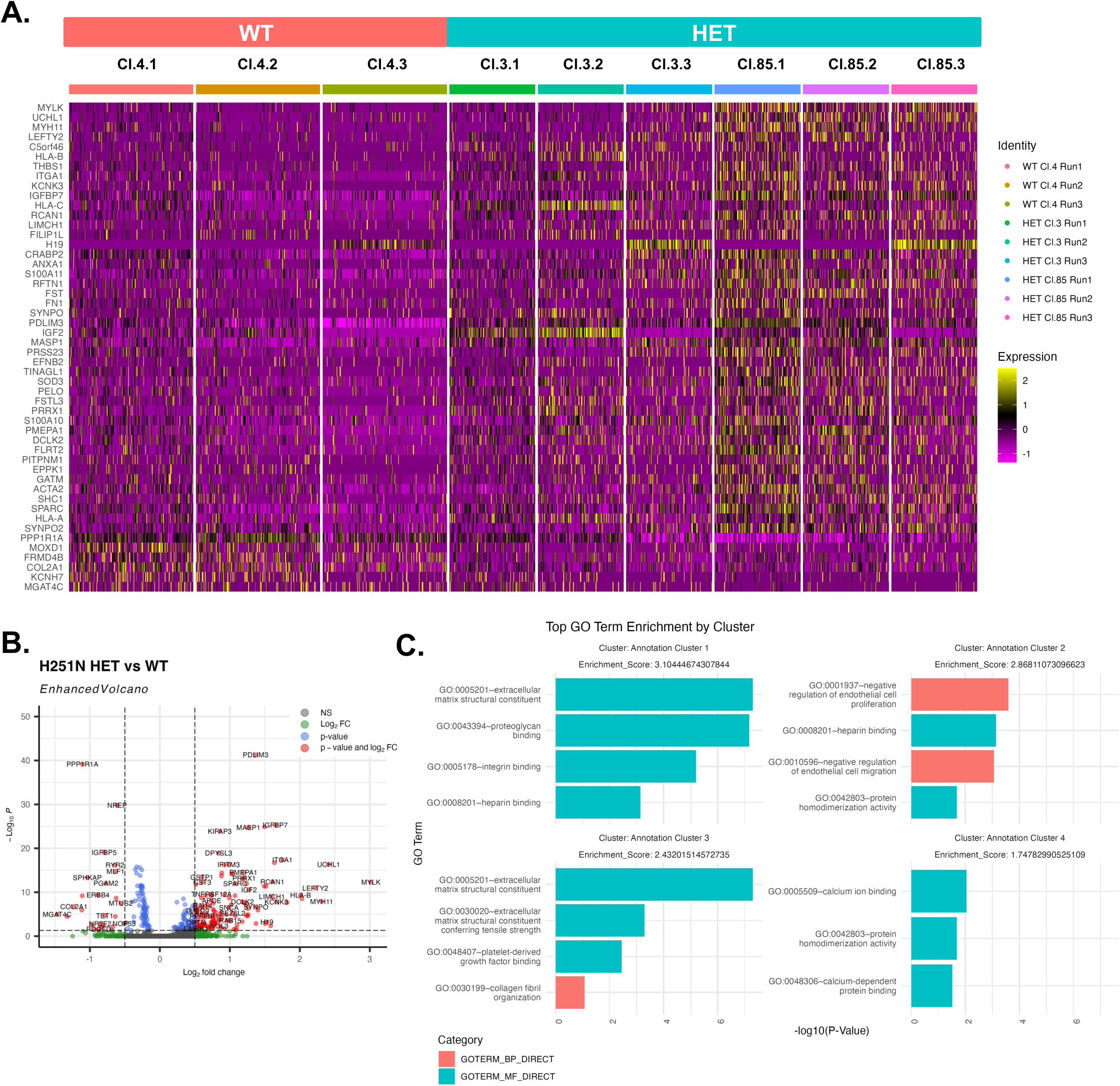

**Supplement Figure 3.**
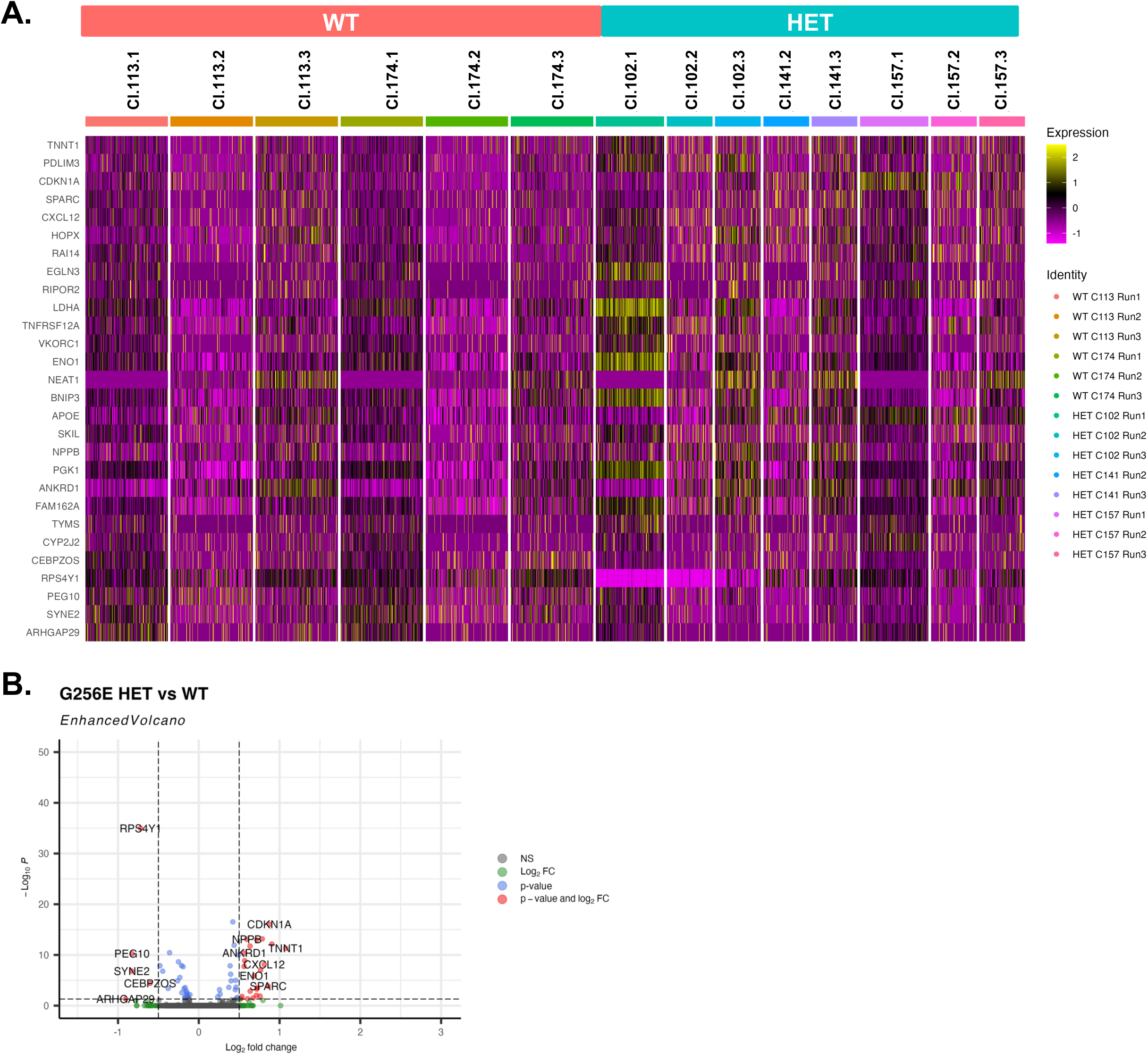

**Supplement Figure 4.**
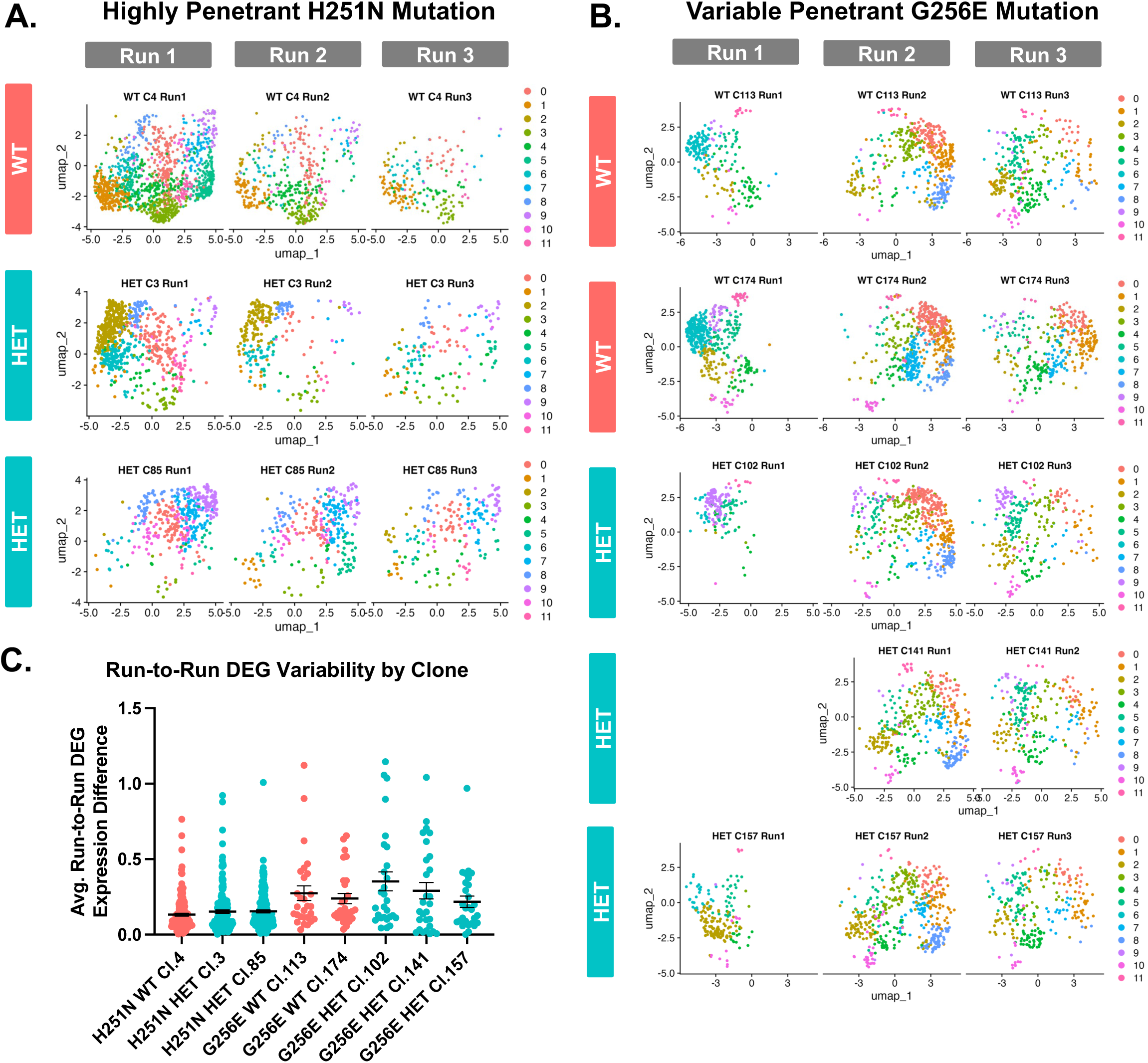

**Supplement Figure 5.**
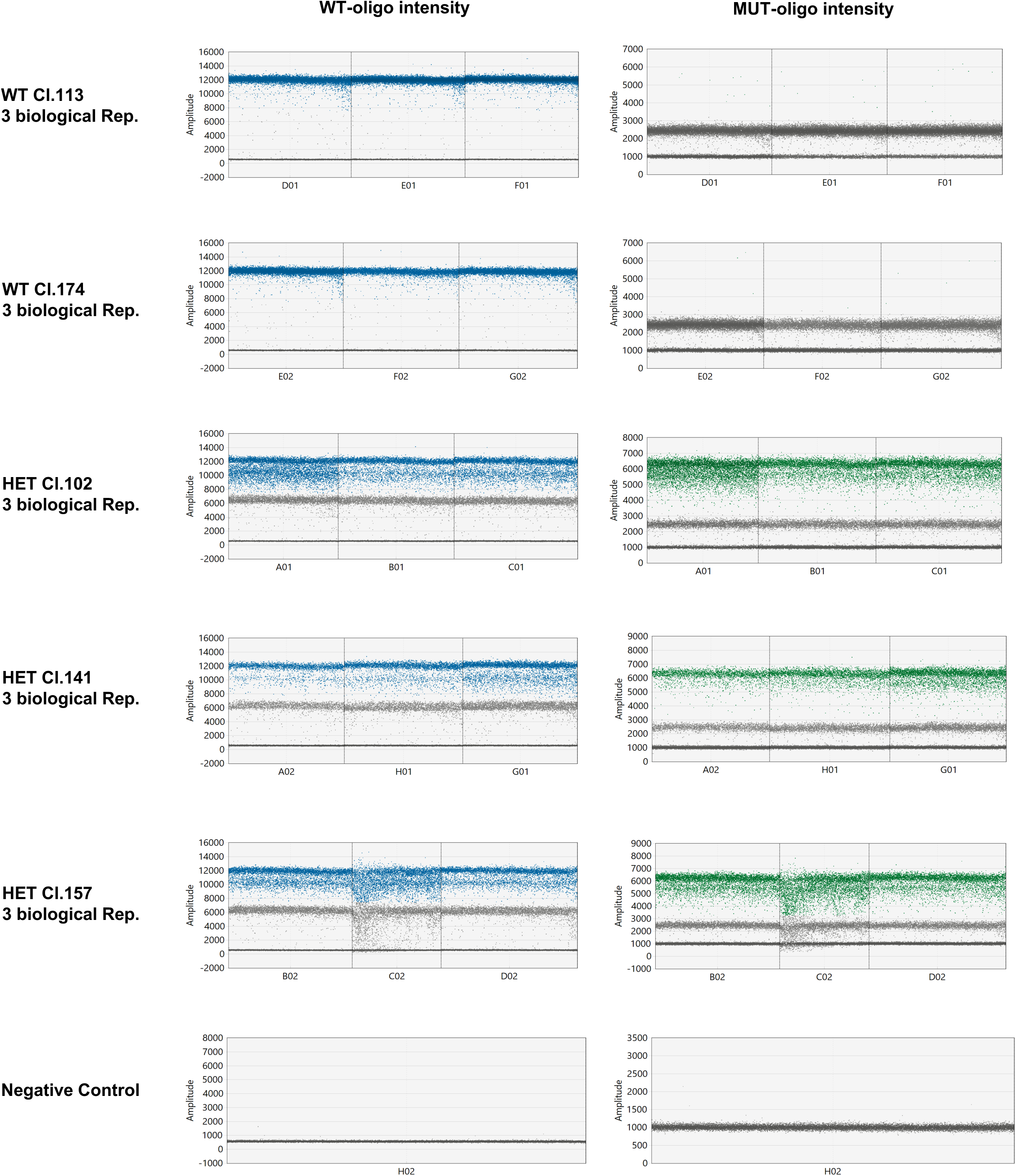

**Supplement Figure 6.**
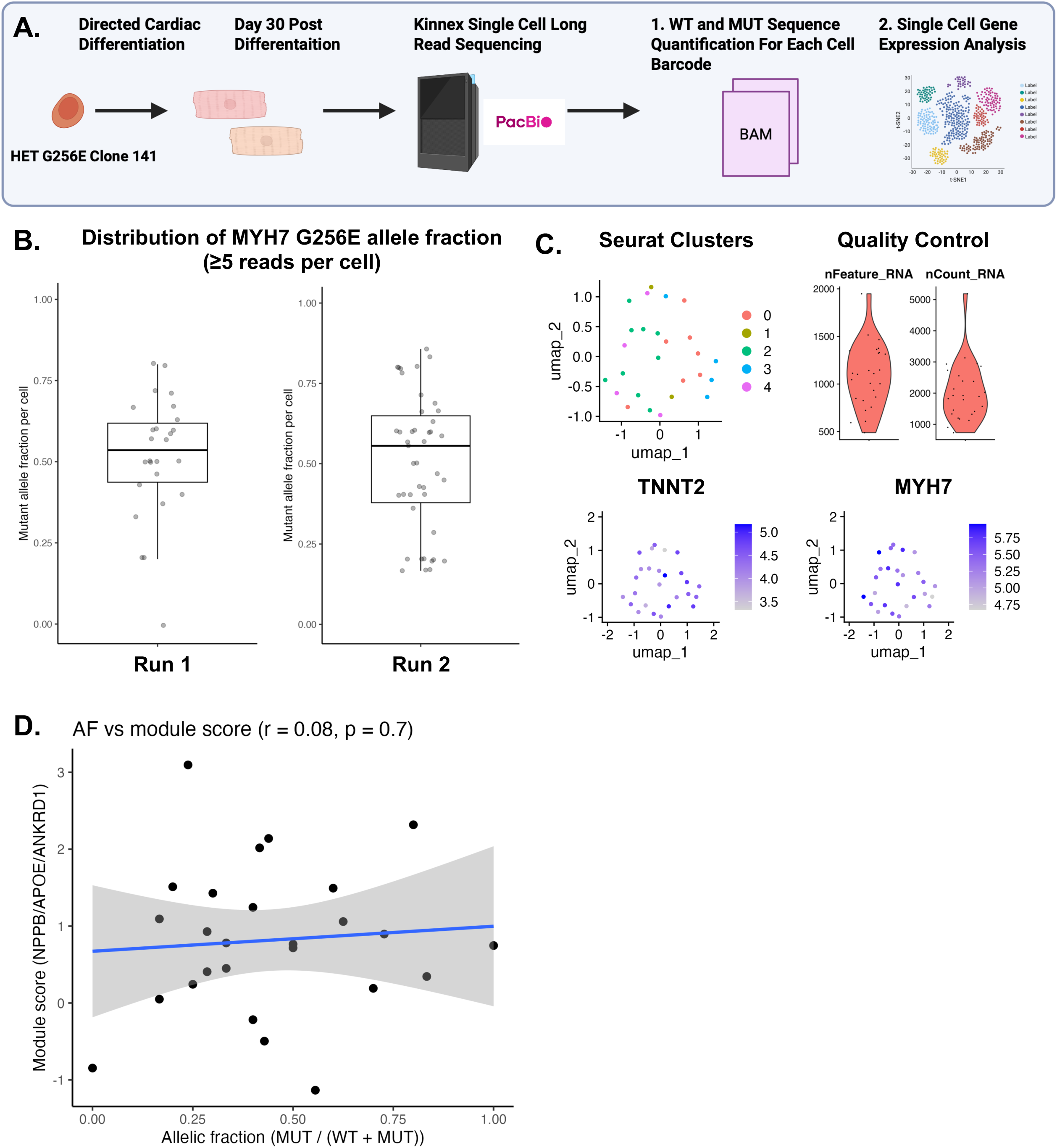

**Supplement Figure 7.**
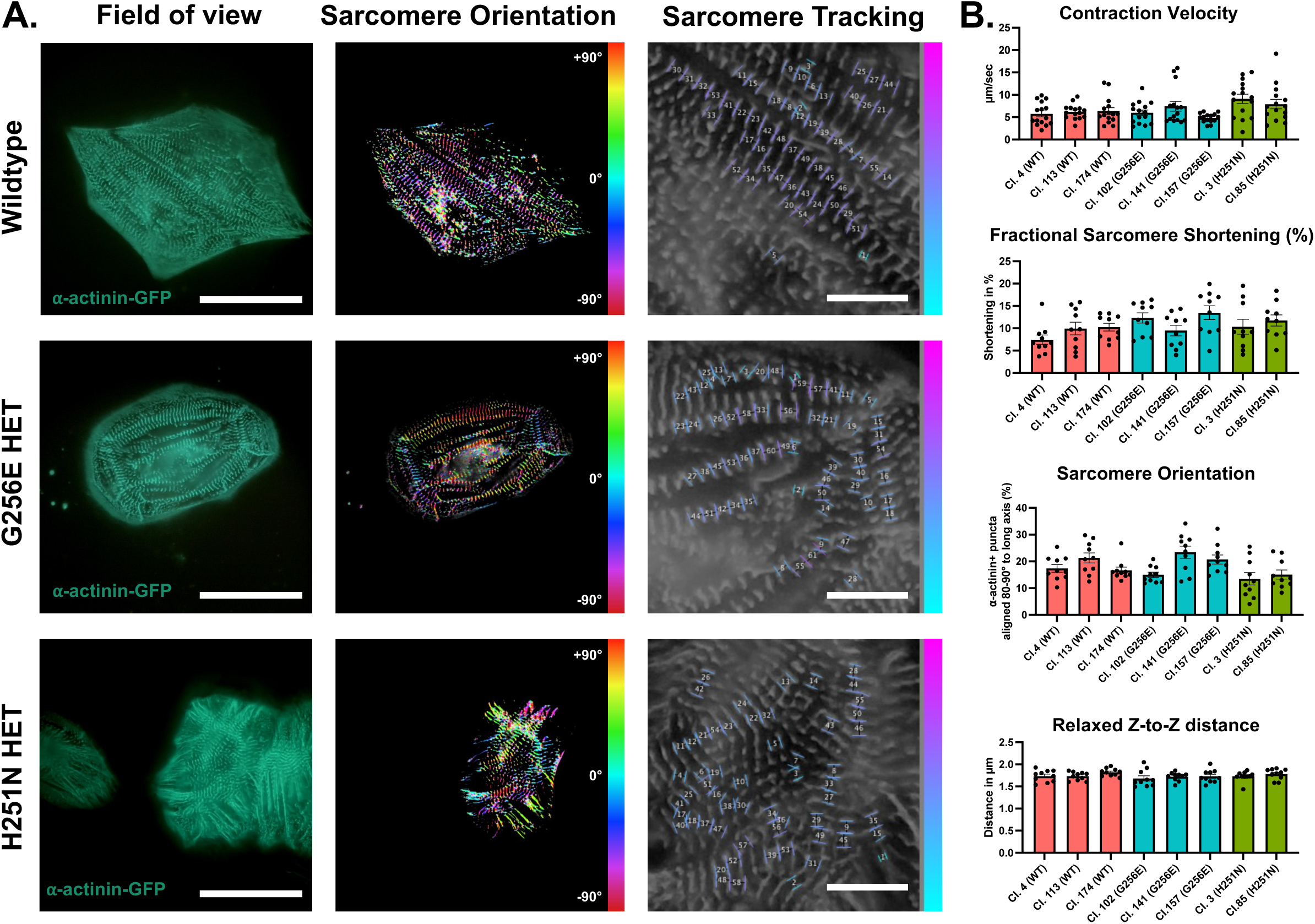

**Supplement Figure 8.**
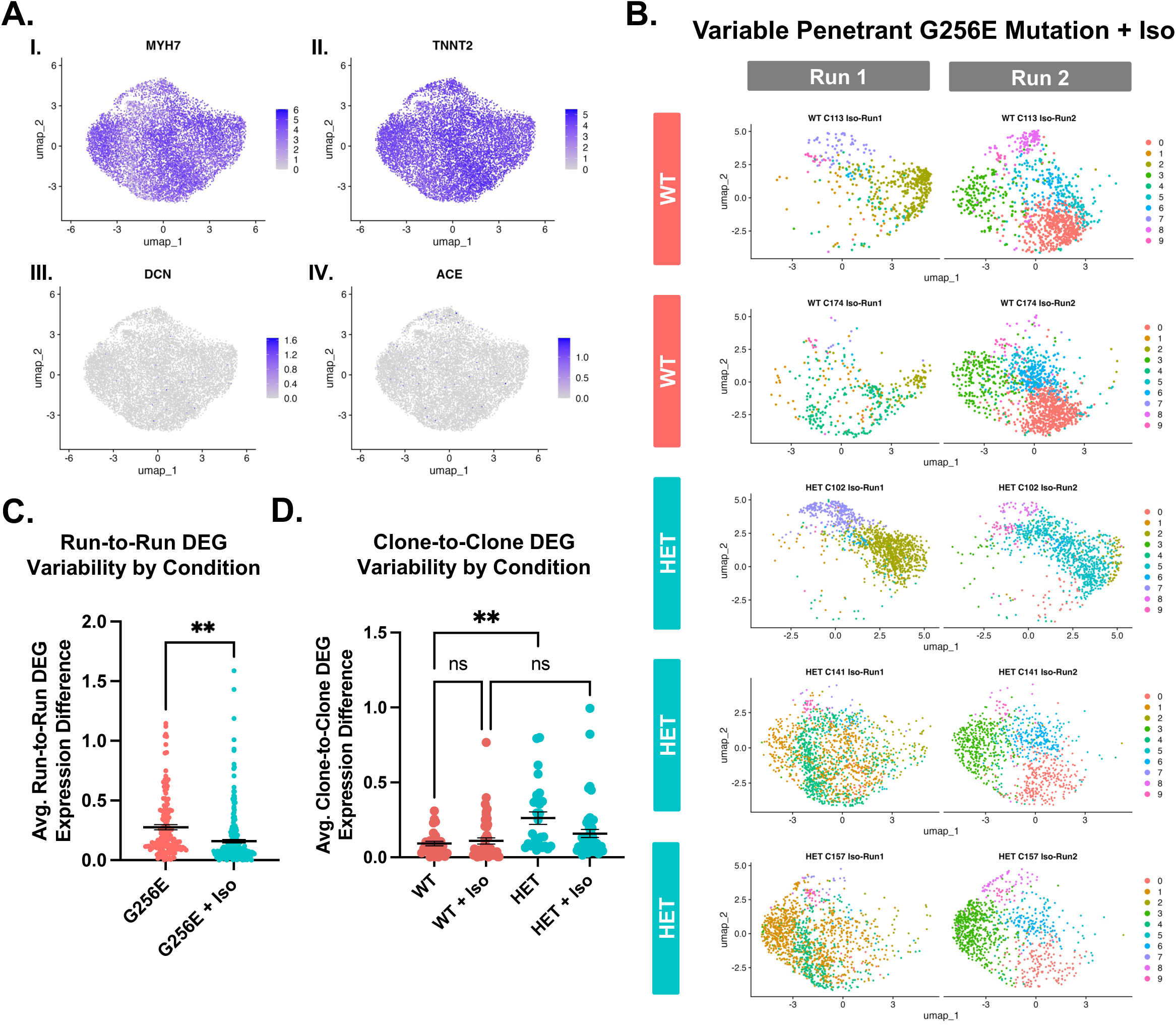

**Supplement Figure 9.**
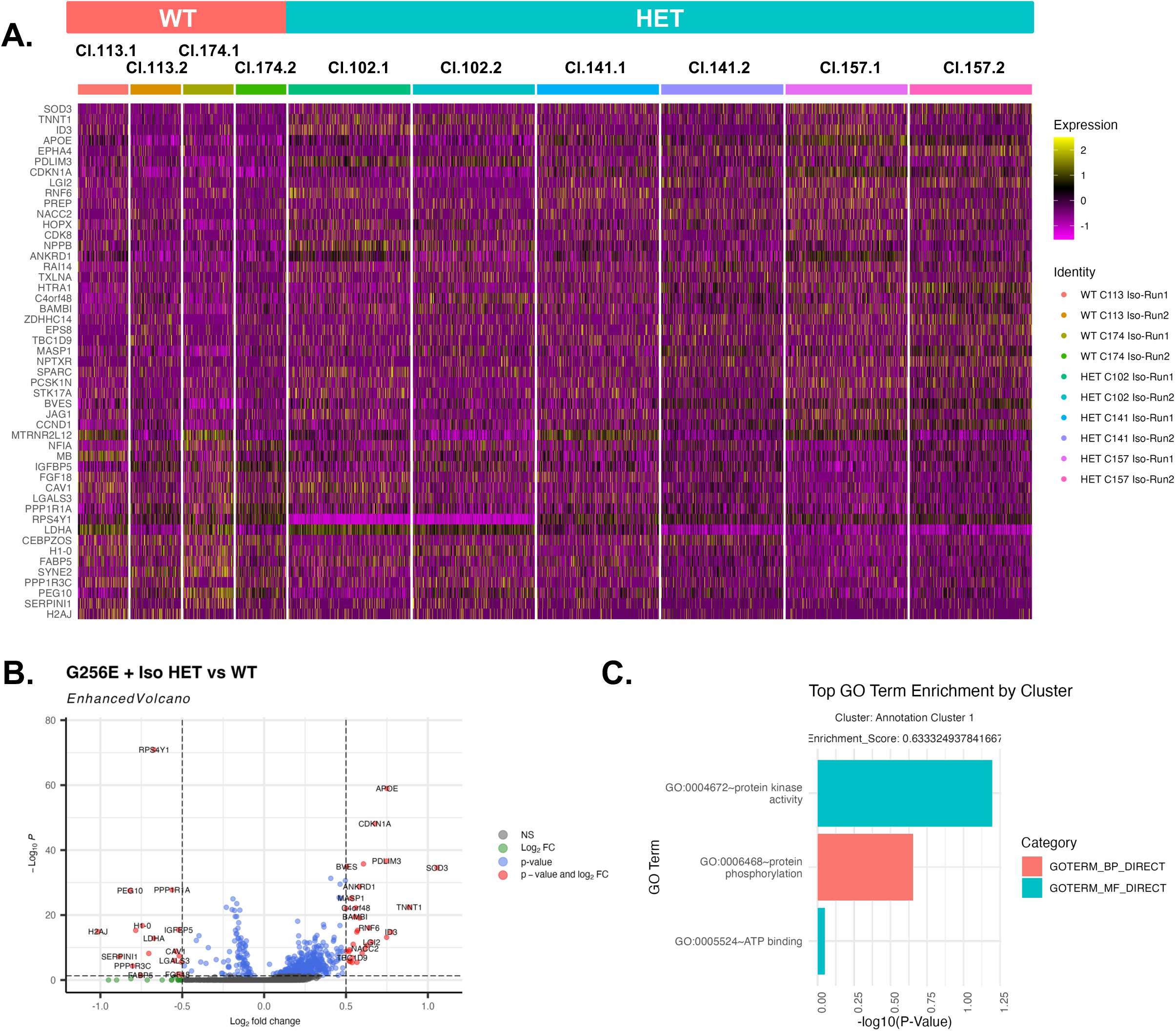

**Supplement Figure 10.**
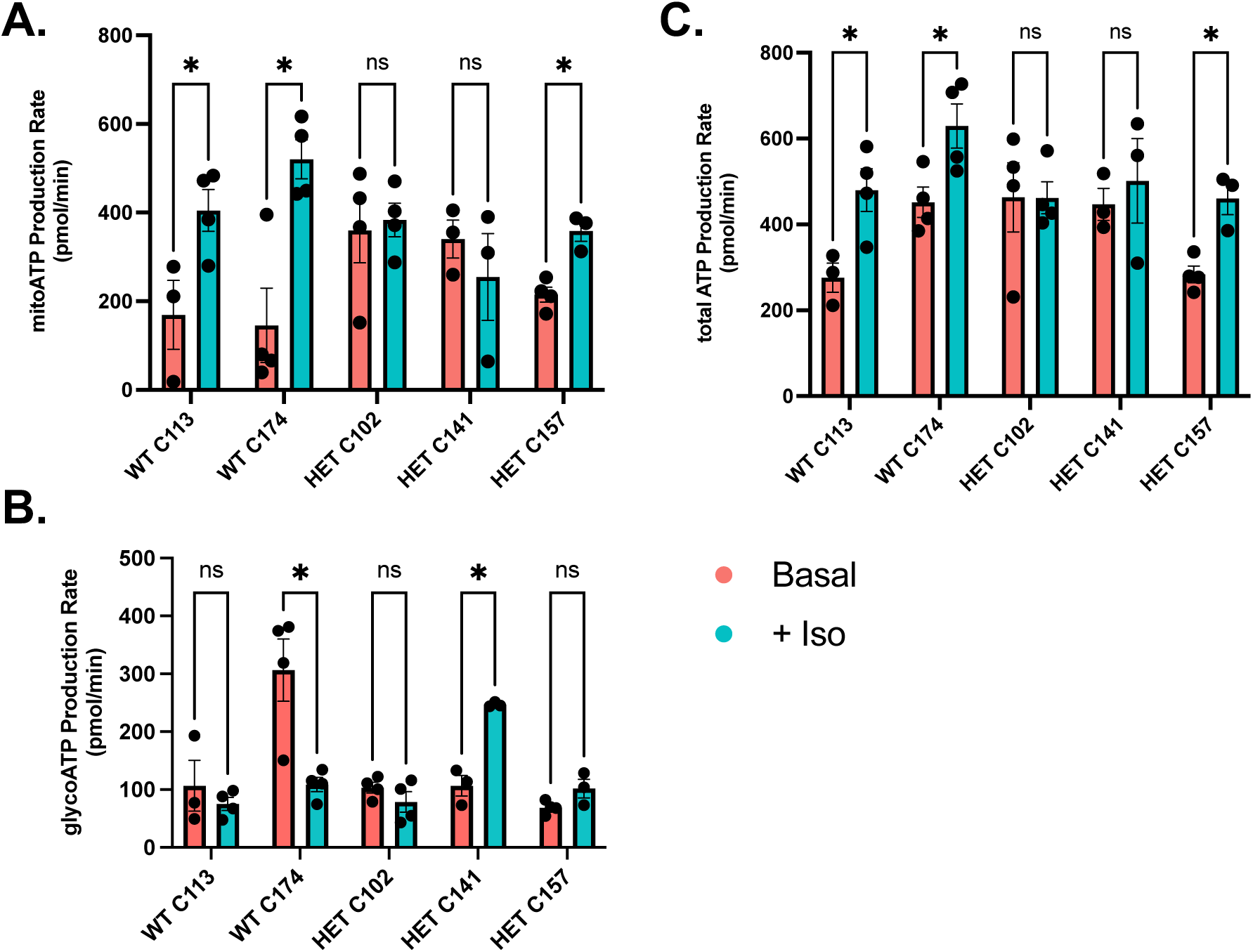

**Supplement Figure 11.**
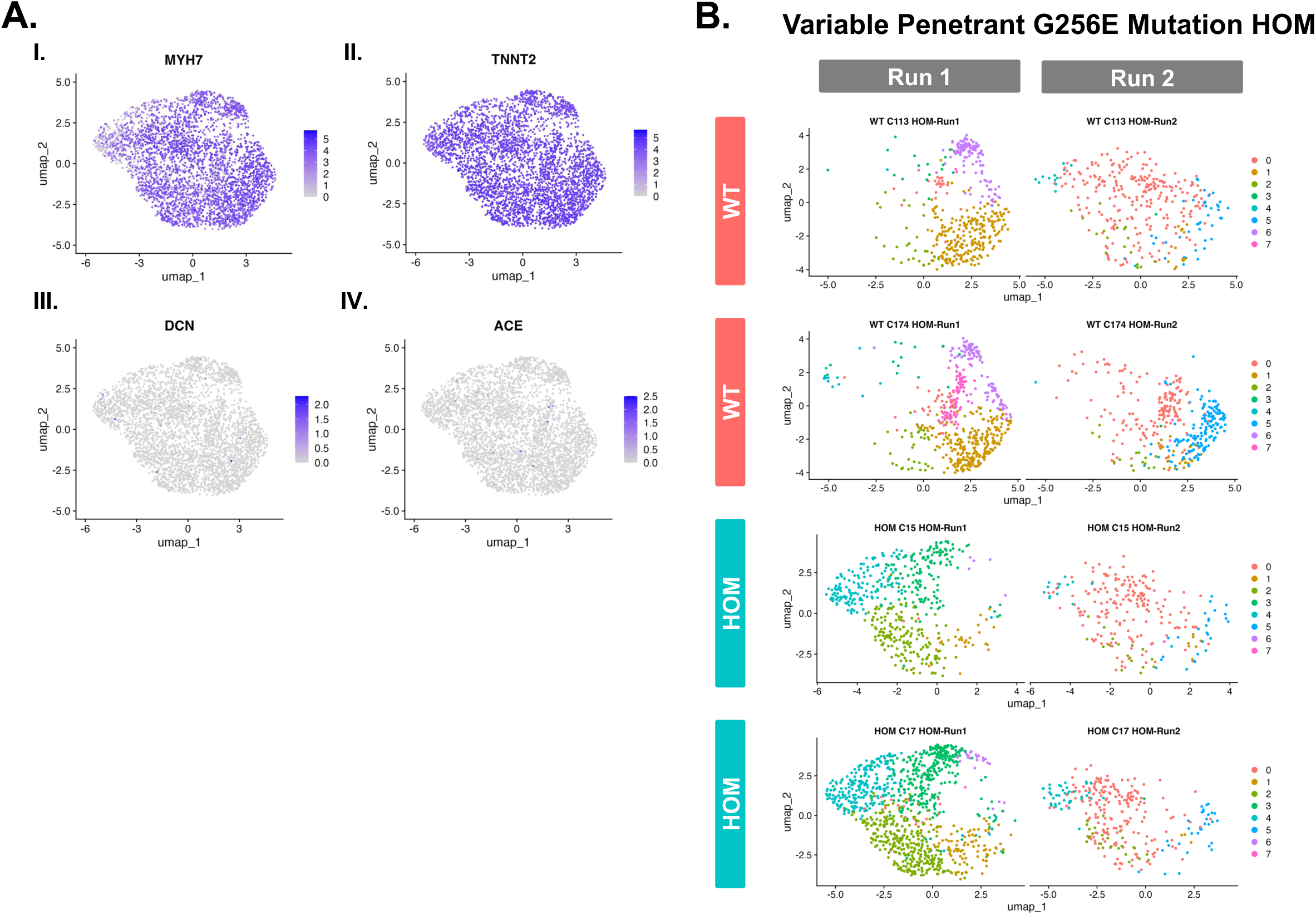

**Supplement Figure 12.**
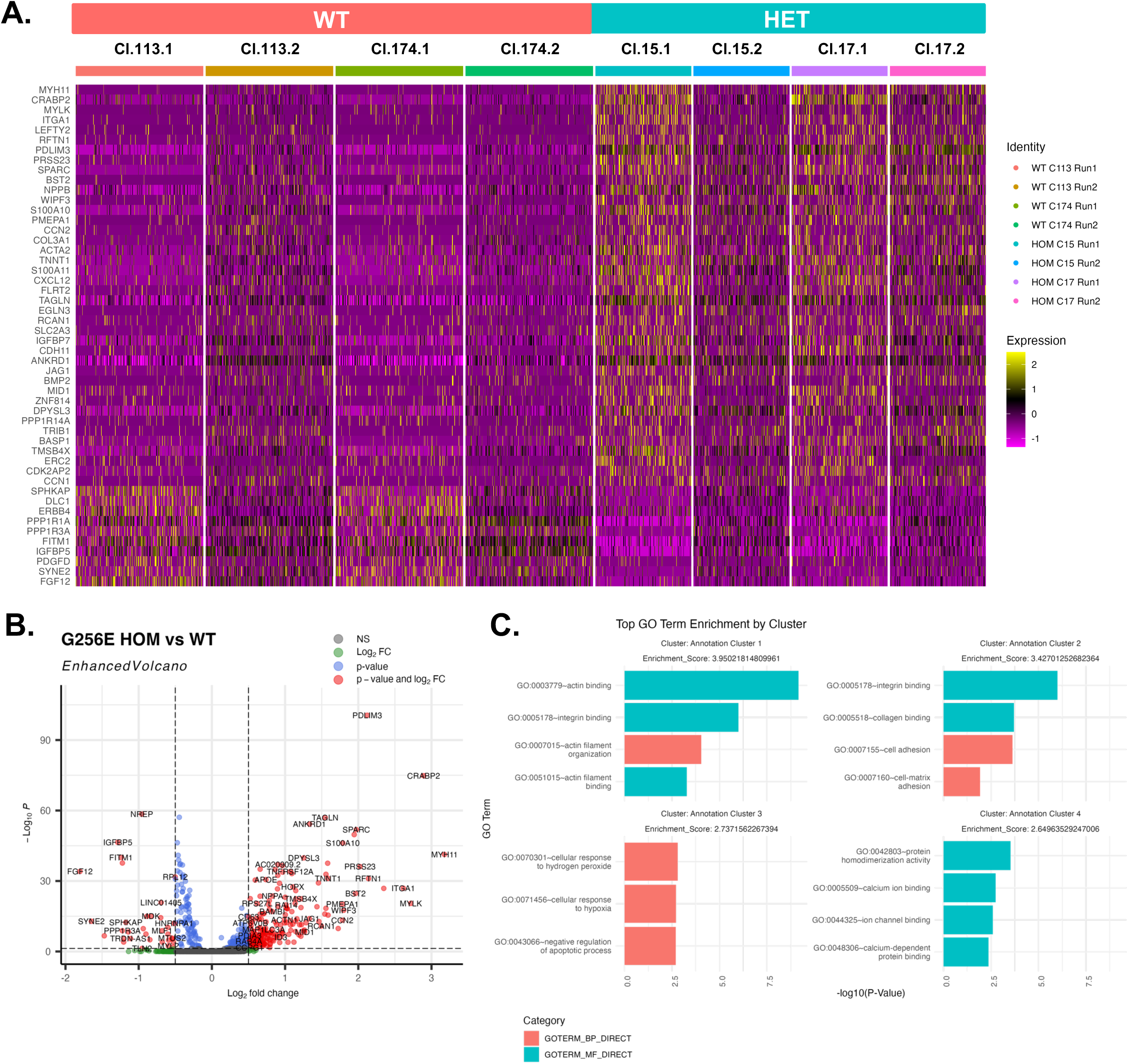

**Supplement Figure 13.**
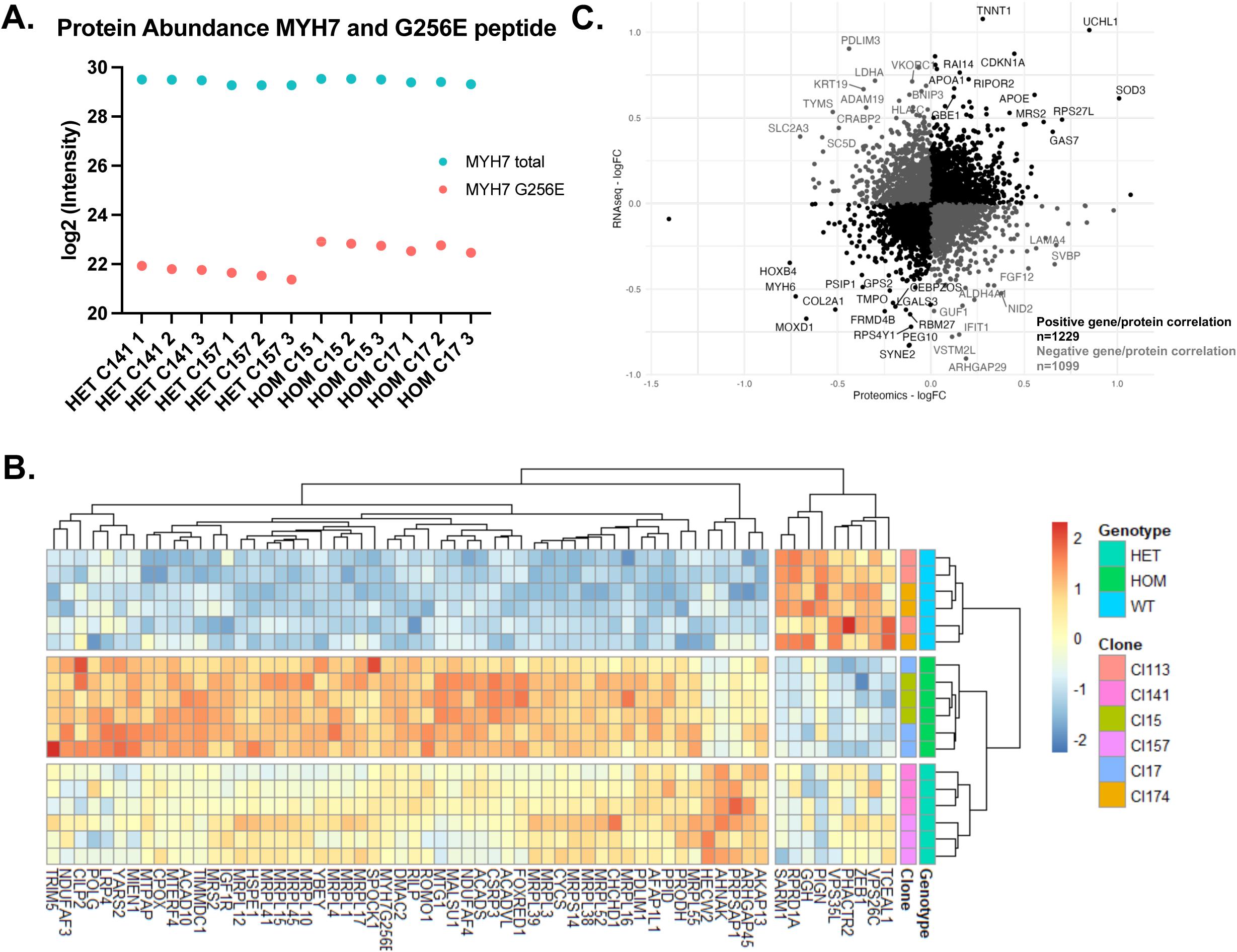

**Supplement Figure 14.**
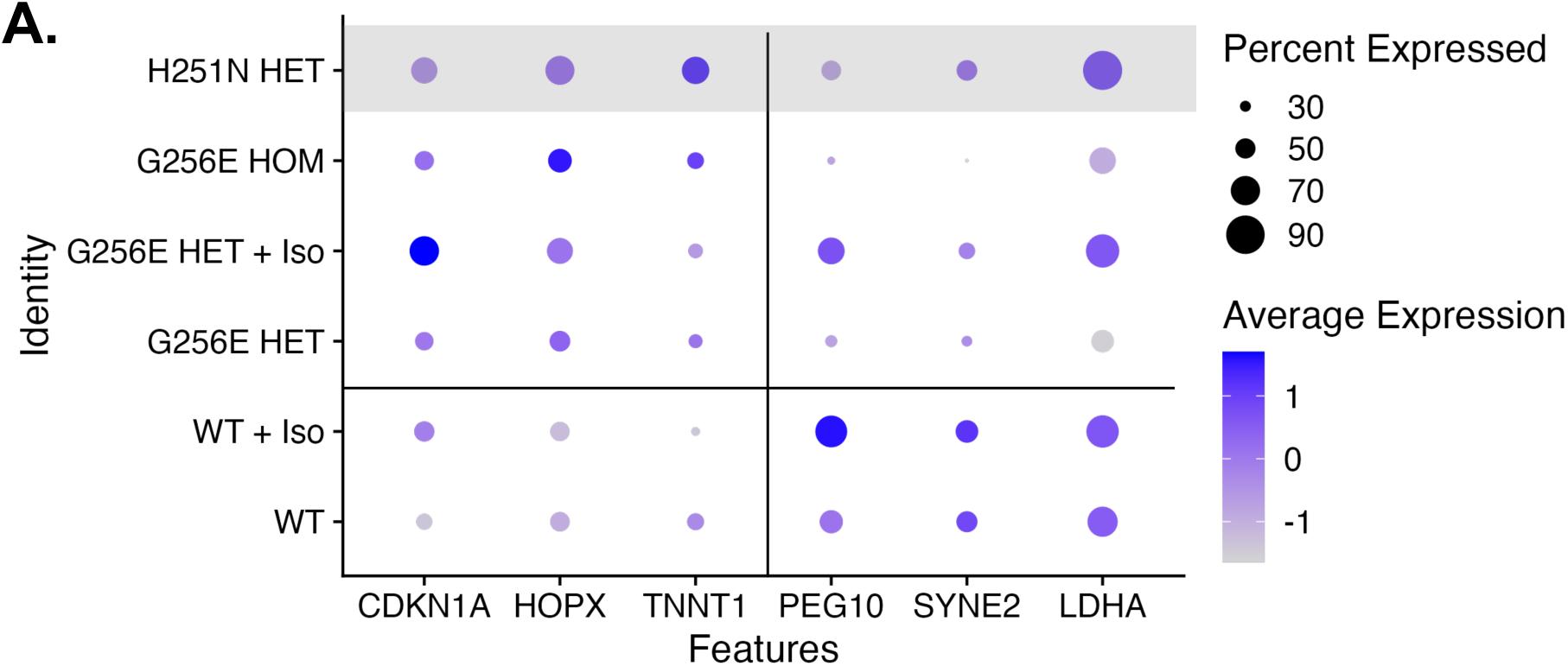

**Supplement Figure 15.**
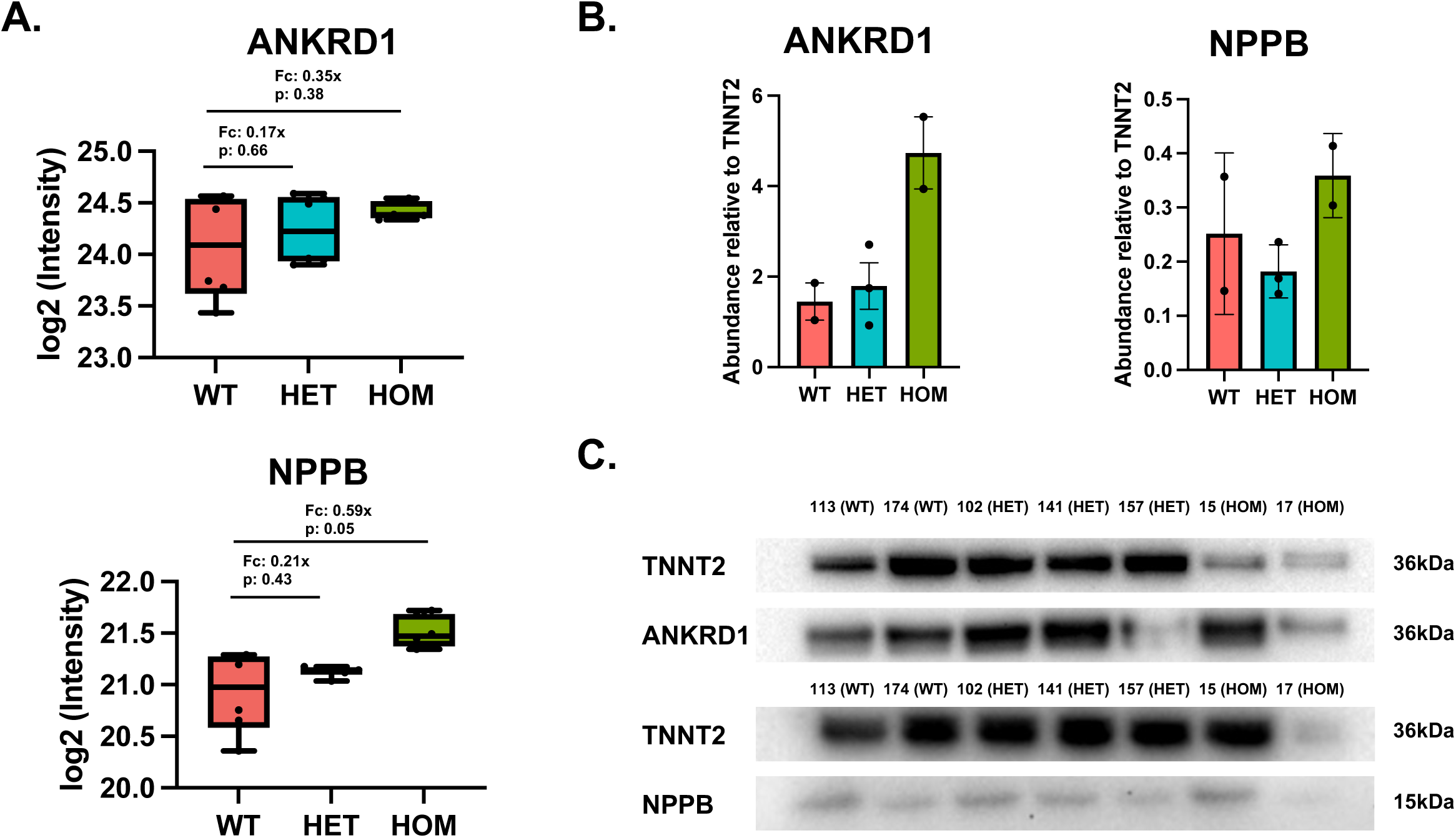

**Supplement Figure 16.**
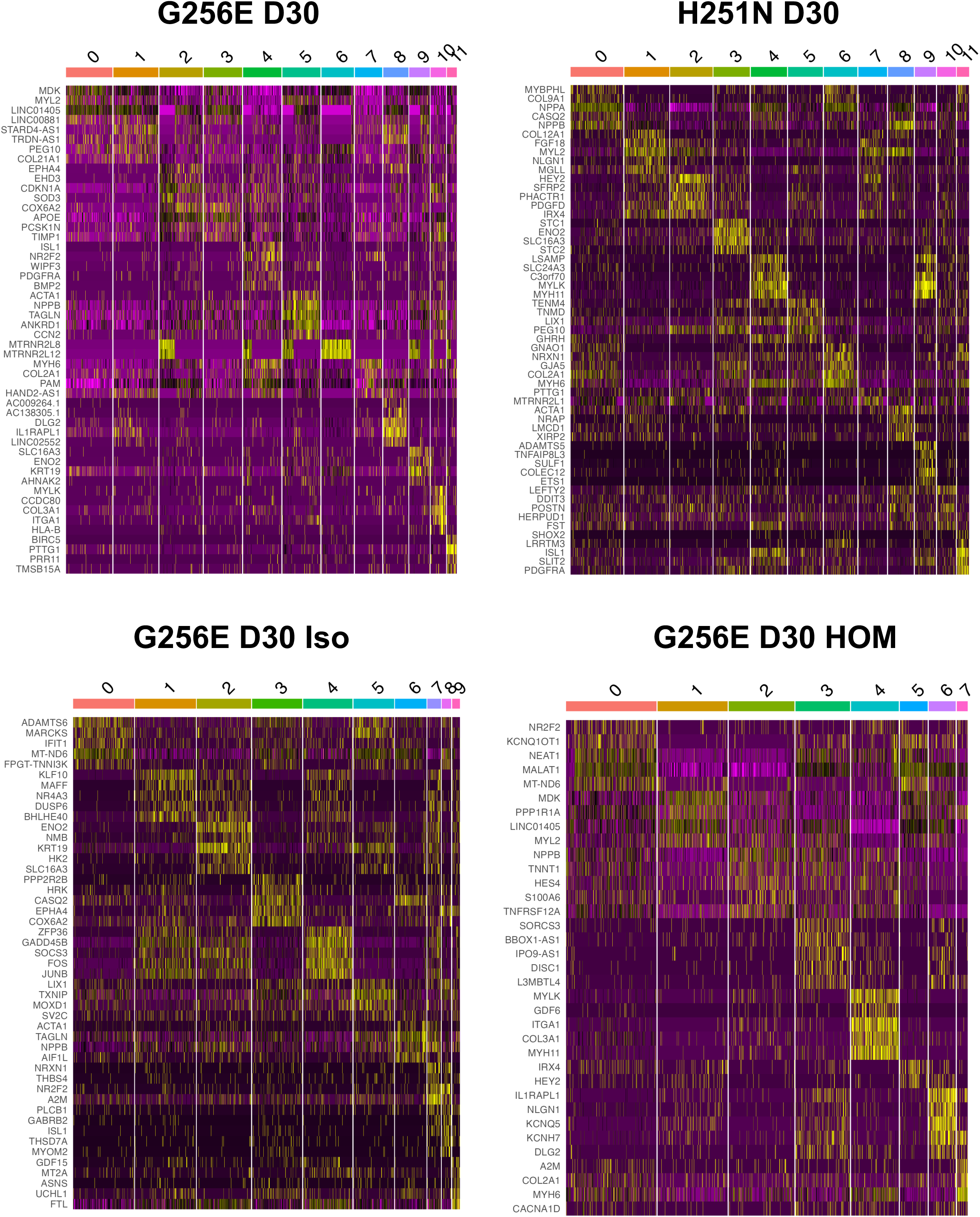

**Supplement Figure 17.**
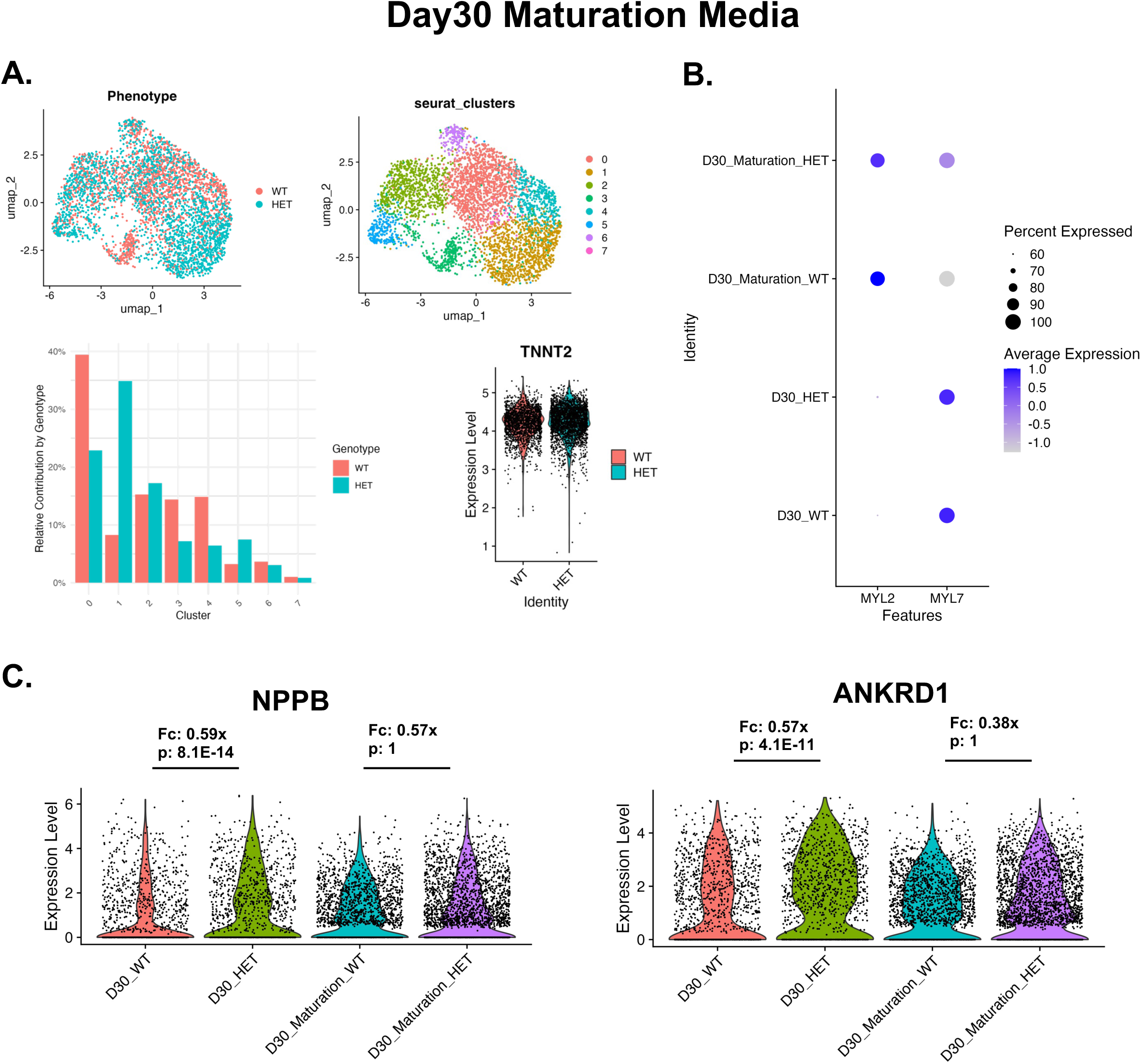

**Supplement Figure 18.**
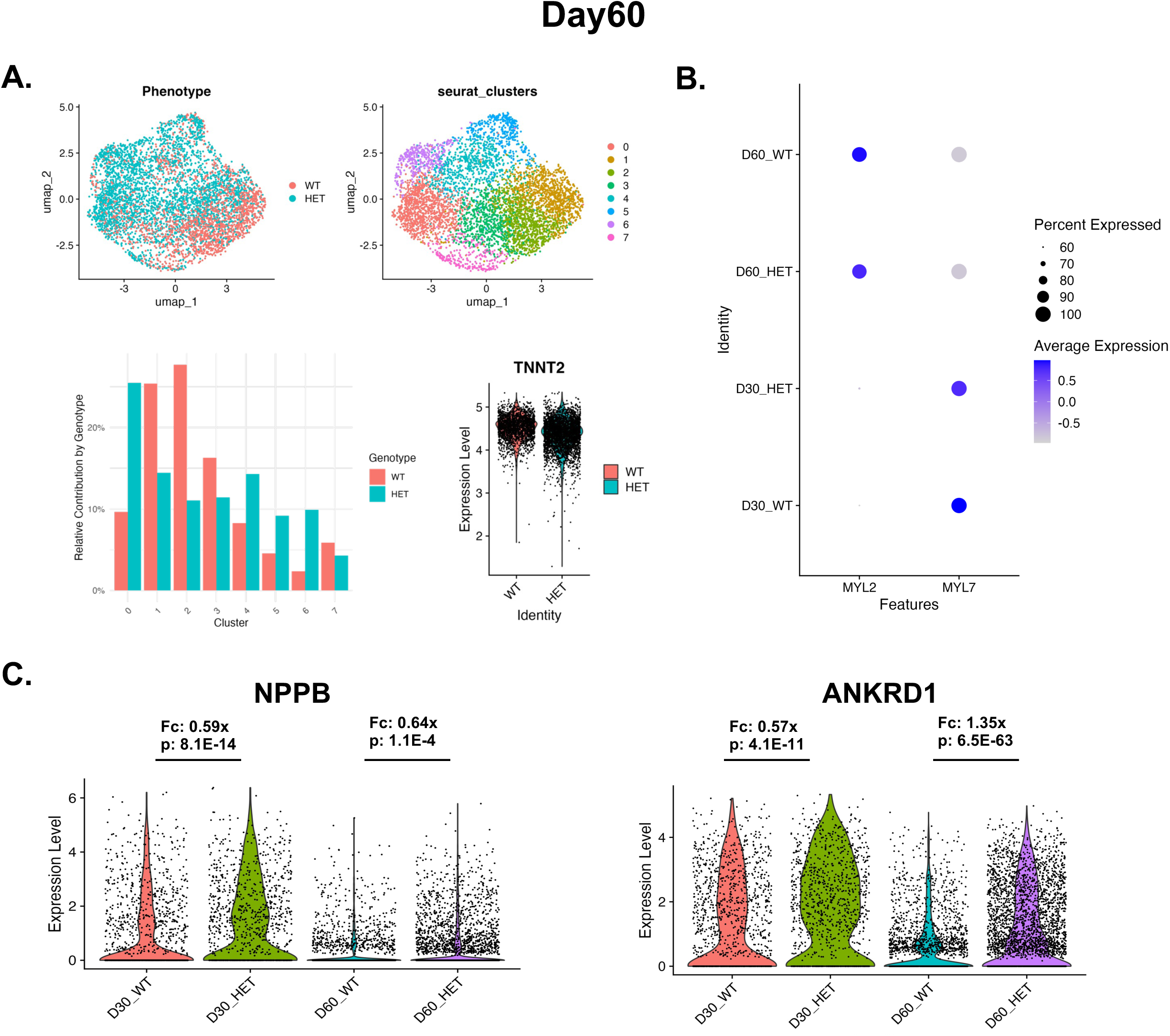

## Notes

### Competing Interest Statement

The authors have declared no competing interest.

