## Supplementary material for "Beta-Adrenergic Stimulation and *MYH7* G256E Mutant Gene Dosage Drive Hypertrophic Cardiomyopathy Phenotype Penetrance": Manuscript_Supplement

**Short Title**: Disease Modeling of Variable Phenotype Penetrance

**Keywords**: Hypertrophic Cardiomyopathy; Variable Disease Penetrance; iPSC-derived Cardiomyocytes

### Supplemental Information (SI)

1. **Extended Methods**
2. **SI Figure Legends**
3. **SI References**

### 1. Extended Methods

*hiPSC culture and CM differentiation*

The parental WTC hiPSC cell line was generated by the Bruce R. Conklin Laboratory at the Gladstone Institutes and University of California-San Francisco (UCSF). Cells were maintained using described methods.[1,2] A detailed cell culture protocol from the Allen Institute for Cell Science can be found on its webpage (http://allencell.org). The parental WTC cell line AICS-0075-085 ACTN2-mEGFP was developed at the Allen Institute for Cell Science (http://allencell.org/cell-catalog) and is available through Coriell (https://www.coriell.org). AICS-0104-004 ACTN2-mEGFP MYH7WT/WT, AICS-0104-003 ACTN2-mEGFP MYH7WT/H251N, AICS-0104-085 ACTN2-mEGFP MYH7WT/H251N, AICS-0097-113 ACTN2-mEGFP MYH7WT/WT, AICS-0097-174 ACTN2-mEGFP MYH7WT/WT, AICS-0097-102 ACTN2-mEGFP MYH7WT/G256E, AICS-0097-141 ACTN2-mEGFP MYH7WT/G256E and AICS-0097-157 ACTN2-mEGFP MYH7WT/G256E were developed at the Allen Institute for Cell Science and are available on request. Homozygous HCM lines WTC ACTN2-mEGFP MYH7 G256E/G256E Cl.15 and WTC ACTN2-mEGFP MYH7 G256E/G256E Cl.17 were derived from AICS-0097-141 ACTN2-mEGFP MYH7WT/G256E via additional gene editing steps described below and are available from the Sean Wu laboratory on request. All cell lines were sequenced and karyotyped to confirm correct gene editing and tested negatively for mycoplasma.

hiPSCs were maintained in a polystyrene 2D culture system coated with Matrigel (Corning) in DMEM/F12 (ThermoFisher) in a 1:400 dilution (coated 24 hours before use). Each hiPSC cell line was maintained in two 6-well plates (Corning) in MTeSR (STEMCELL Technologies) hiPSC serum free maintenance media. One well was passaged in a 1:12 splitting ratio upon 80-85% confluency using StemPro Accutase(Gibco) for single-cell-dissociation. The hiPSCs in the remaining 11 wells were differentiated into CMs through directed cardiac differentiation. For passaging, 10uM of ROCK inhibitor Y-27632 was added to the MTeSR maintenance media for the first 24 hr after replating. hiPSC-CM differentiation was initiated using the canonical Wnt stimulation and inhibition protocol[3] in serum free RPMI 1640(Invitrogen)-based differentiation media supplemented with B27 minus insulin (Invitrogen). A gradient of CHIR99021 (Selleckchem) concentrations (7.0, 7.5, 8.0 μM) was used between days 0-3 of differentiation. Between days 3-5, Wnt-C59 (2.0 μM) (Selleckchem) was added to the differentiation media, and between days 7-9, RPMI 1640 was supplemented with B27 with insulin. Between days 9-11, RPMI 1640 minus glucose was supplemented with B27 with insulin, and on day 11, RPMI 1640 with B27 with insulin was used as a maintenance media. On day 12, wells that contained more than 90% of the beating CMs were treated with TrypLE Select Enzyme 10X (Invitrogen) at 37 °C for 5-10 minutes. RPMI

1640 + B27 with 10% Knock Out Serum Replacement (Gibco) and Thiazovivin 1.0 μM (Selleckchem) was added in a 1:1 ratio, and the cells dissociated from the well were then transferred to a 50 mL conical tube and centrifuged at 1000 RPM for 3 minutes. Afterwards, the pellet was resuspended in 1mL BamBanker (ThermoFisher) freezing media per well, and 1mL of the resuspended pellet was aliquoted into individual cryogenic vials. The cryogenic vials were stored at -80°C overnight and transferred to a liquid nitrogen tank the next day for long-term storage. For downstream experiments, frozen day 12 CMs were thawed using a 37 °C metal-bead bath and resuspended in replating media RPMI 1640 + B27 with 10% Knock Out Serum Replacement (Gibco) and Thiazovivin 1.0 μM (Selleckchem). Cell suspension was spun down at 1000 RPM for 3 minutes. CMs were resuspended in 2ml replating media and plated on to one 6 well. Media was changed to RPMI 1640 with B27 with insulin the next day and changed every other day for CM maintenance.

*Droplet-based single cell RNA sequencing*

One batch of day 30 hiPSC-CMs per cell line were collected and individually labeled using 10X Genomics CellPlex cell labeling technology. First, hiPSC-CMs were dissociated into single cells using TrypLE 10X Select (Gibco). The cells were washed three times with replating medium containing 10% knock-out serum replacement, 1μM thiazovivin, and RPMI-1640 supplemented with B27+ insulin supplement and filtered to remove any cell aggregates. They were then resuspended in PBS with 0.04% BSA at room temperature. CMs were counted and 500,000 cells per cell line or experimental condition were centrifugated and labeled by resuspension in 100 μl Cell Multiplexing Oligo (CMO). CMOs and cells were co-incubated for 5min at room temperature. After incubation, the cells were washed three times with PBS containing 1% BSA to remove unbound CMOs at 4 °C. CMs were then counted, and equal amounts of each sample were pooled for downstream processing. The labeled single-cell samples were loaded into the Chromium Controller Instrument (10X Genomics) to generate single-cell-barcoded droplets (GEMs) using, following the manufacturer's protocol. The resulting libraries were sequenced on an Illumina NovaSeq instrument using the 10X recommended read length. At least two independent scRNAseq runs were conducted per experimental condition, and the data were bioinformatically integrated as described below. The data are publicly available under GEO accession number GSE276210.

*ScRNAseq Data Pre-Processing*

10X Cell Ranger Multi v7.2.0, a set of analysis pipelines that process Chromium single cell data, was used to pre-process scRNAseq data including sample demultiplexing, alignment, filtering, barcode processing, and gene counting. FASTQ files were generated by Illumina's bcl2fastq. Then, reads in FASTQ files were mapped to the human reference genome (GRCh38 2020-A).

*ScRNAseq Data Analysis*

For the downstream data analysis, Seurat R package was used.[4] First, cells with <200 or >10.000 detected genes and <250 or >150.000 RNA counts were filtered. Cells with a percentage of mitochondrial genes >50% were additionally filtered. The raw read counts generated by 10X

CellRanger were subsequently normalized using a global scaling method called "LogNormalize." This method involves normalizing the gene expression measurements for each cell based on the total expression, scaling, and then applying a log transformation to the results. The cells were then placed in low-dimensional space using non-linear dimensional reduction analysis such as Uniform Manifold Approximation and Projection (UMAP). Next, graph-based unbiased clustering was performed using a clustering resolution parameter of 1.2 for baseline conditions in Figure 1 and 0.8 in Figure 2 and 3. Cells with normalized counts were further assigned to a cell cycle state using cell-cycle scoring and regression function “Cell Cycle Scoring” in Seurat and cells in G1 cell cycle phase were subsetted for further analysis. Subsequently, *TNNT2* expression was calculated per cell cluster and non-myocyte clusters (defined by normalized *TNNT2* expression value of <30) were removed. Finally, subsetted cells were re-embedded and clustered and each cluster was assessed for quality control (QC) markers gene count per cell, RNA count per cell and proportion of mitochondrial gene expression. If final clusters showed QC marker values below two standard deviations of all clusters, they were removed. Different scRNAseq batch data were integrated using the Harmony method, which corrects for batch effects by iteratively adjusting the data to align shared gene expression patterns while preserving biological variability.[5] To quantify the contribution to each cluster by underlying genotype of the cell, we calculated the counts of each genotype within each cluster and then divided it by the summed the counts for each genotype across all clusters. Differentially expressed genes (DEGs) between mutant cells and the isogenic controls were determined by detecting genes, where the expression fold-change is >1.4 (logFC > 0.5) and adjusted p-value is < 0.05 with at least 20% of cells expressing the identified gene in one of the experimental groups. To account for different amounts of single cells in each experimental group and to avoid batch specific gene expression as a confounding factor in the DEG analysis, cells from each clone and individual batch within an experimental group were down sampled to the same number of single cells per group prior to DEG analysis. The log-transformed normalized single cell gene expression values were used for visualizations (violin plots) and differential expression tests. The log-transformed normalized scaled single cell gene expression values were used for DEG visualization (heatmap). For gene ontology (GO) analysis, the Database for Annotation, Visualization, and Integrated Discovery (DAVID) tool was used to identify significantly altered biological processes. Briefly, the upregulated and downregulated DEGs were submitted to DAVID Bioinformatics Resources Database (https://david.ncifcrf.gov/home.jsp) for identifying enriched GO-biological process and GO-molecular function terms. Functional annotation clustering was then used to identify functional groups within the identified terms. To assess run-to-run variability, we extracted relevant data, including clone, run, and markers of interest (DEGs), using the FetchData function and converted it to a data frame. For each clone we calculated the average expression levels of previously identified DEGs for each individual run. Absolute differences between these averages were computed to determine run-to-run expression variability for each clone. To assess clone-to-clone variability, we extracted relevant data, including clone, genotype and markers of interest (DEGs), using the FetchData function and converted it to a data frame. Within each genotype, we then calculated the average DEG expression difference between single clones of this specific genotype to determine clone-to-clone DEG expression variability.

*Digital Droplet PCR*

Primers were designed to specifically amplify a region of the human *MYH7* gene around the G256E mutation site. Primers were validated for *MYH7* specificity in silico with NCBI's primer designing tool Primer-BLAST (https://www.ncbi.nlm.nih.gov/tools/primer-blast/index.cgi). Two hydrolysis probes with different fluorophores (HEX & FAM) were used to detect either the wild-type or the MYH7 G256E mutant allele in a single reaction with the above primers. Primers and probes were ordered from IDT (Coralville, IA); probes contained the fluorophore at the 5′ end, the IowaBlack quencher at the 3′ end, and an additional internal ZEN quencher. All oligos were stored in the dark at 4 °C for less than 1 year in TE buffer. Primer/probe type, name, reaction concentration, and sequence are listed below:

- Forward; MYH7-mRNA-F/1; 1.8 μM; CCGCTTCGGGAAATTCATTCG
- Reverse; MYH7-mRNA-R/1; 1.8 μM; TGCTTTCAGCTGGAAAATAACTCT
- Wild-type probe; MYH7-WT-mRNA-P/1; 0.25 μM; FAM- aacagGaaagttggcatctgcag-Zen/IowaBlack
- Wildtype probe; MYH7-MU-mRNA-P/1; 0.25 μM; HEX- aacagAaaagttggcatctgcag-Zen/IowaBlack

Each ddPCR reaction was prepared and analyzed with the QX200 ddPCR system (BioRad, Hercules, CA) per BioRad's standard recommendations for use with their ddPCR™ Supermix for Probes (No dUTP). All reactions were mixed to 25 μl and contained up to 10 μl of cDNA reverse transcribed RNA lysates of wildtype and heterozygous G256E CMs. For droplet generation, 20 μl were loaded into the droplet generator cassette in groups of eight per BioRad's protocol. Thermocycler conditions: 95 °C × 10 min; 50 cycles of 94 °C × 30 s and 60 °C × 60 s; 98 °C × 10 min; hold at 4 °C. Data analysis was performed with QuantaSoft Analysis Pro (BioRad, Hercules, CA).

*Long-Read Sequencing*

Allele-specific single-cell transcript quantification was performed using a long-read sequencing approach. HET *MYH7* G256E hiPSC-CMs (day 30) were processed using the 10x Genomics 5′ single-cell capture platform. Full-length cDNA libraries were generated using the PacBio Kinnex Iso-Seq workflow and sequenced on the PacBio Revio system. Long-read sequences from raw BAM-files were used to distinguish WT and G256E mutant *MYH7* transcripts (69 cells after QC), and reads were assigned to individual cells based on 10x cellular barcodes (26 cells after QC). Allelic fractions were calculated per cell as the proportion of mutant versus mutant+WT *MYH7* reads. In parallel, standard 10x workflow was used for transcriptome analysis and UMAP generation.

*Contractility Measurement and Structural Analysis*

The contraction velocity of single hiPSC-CMs was measured on day 35 of differentiation using a cell motion imaging system (Sony SI8000, Sony Biotechnology). CMs were sparsely replated on day 27 at a density of approximately 40,000 cells/cm². On day 35, spontaneously beating individual cells were selected, and videos were recorded using a 10× objective for 8 seconds at

150 frames per second (fps) with a resolution of 1024 × 1024 pixels. Contraction velocity was quantified using edge-detection–based analysis implemented in the Sony SI8000 software. For sarcomere-based analysis, CMs were replated on day 27 onto 35 mm glass-bottom dishes at a density of approximately 40,000 cells/cm² and imaged on day 35. All cell lines used in this study carry an α-actinin–GFP (ACTN2-mEGFP) reporter, which was used to visualize Z-disks. Imaging was performed on a Zeiss LSM980 microscope, capturing the native GFP fluorescence. 10s videos were acquired at 30 fps using a 63× objective for live Z-disk motion tracking. Sarcomere orientation was quantified using OrientationJ (ImageJ)[6] by measuring the proportion of α-actinin–positive structures aligned at 80–90° relative to the long axis of the cell. Fractional sarcomere shortening and relaxed Z-to-Z distance were analyzed using SarcTrack implemented in MATLAB.

*β-Adrenergic stressor treatment of Day 30 hiPSC-CMs*

2M frozen day 12 hiPSC-CMs were thawed then transferred to a 15mL conical tube containing 5mL of replating media RPMI 1640 + B27 with 10% Knock Out Serum Replacement and Thiazovivin 1.0 μM and centrifuged at 1000 RPM for 3 minutes. The pellet was resuspended with replating media and transferred to one well of a 6-well plate. The next day, the replating media was replaced with RPMI 1640 supplemented with B27 with insulin and replaced every other day. On day 27, the hiPSC-CMs were replated to a 48-well plate at 40.000 cells/cm$^2$ for contractility measurement and to a 12well plate at 160.000 cells/cm$^2$ for scRNAseq downstream analysis. Starting on day 30, the hiPSC-CMs were treated with 1μM isoproterenol (Sigma Aldrich) in RPMI 1640 supplemented with B27 with insulin for five days with media changes every day.

*Analysis of Cell Metabolism (Seahorse)*

Seahorse ATP Rate Assay (Agilent, USA) was performed according to the manufacturer's instruction. Briefly, 50,000 hiPSC-CMs were plated into Matrigel coated 96-well Seahorse plates. Following Isoproterenol treatment (1 μM) for 5 days, the oxygen consumption rate (OCR) and extracellular acidification rate (ECAR) were measured. One hour before the assay, culture medium was exchanged with Agilent Seahorse XF RPMI Basal media supplemented with 2 mM glutamine, 10 mM glucose and 1 mM pyruvate. Electron transport chain uncouplers were diluted in the same media and sequentially injected during the measurements at the following final concentrations; Isoproterenol (1 μM), Oligomycin (2.5 μM), and Rotenone and antimycin A (1 μM, each).

*Generation of G256E HOM Clones*

A specific single-stranded oligonucleotide (ssODN) was designed uniquely for the G256E mutation located in the *MYH7* gene within the ACTN2-mEGFP cell line and ordered from IDT: 5' AATTCATTCGAATTCATTTTGGGGCAACAGAAAAGTTGGCATCTGCAGACATAGAGACC 3'. Custom synthetic sgRNA targeting the protospacer sequence of 5' TTTTGGGGCAACAGGAAAGT 3' were ordered also from IDT. Recombinant wild-type Streptococcus pyogenes Alt-R® S.p. Cas9 Nuclease V3 was purchased from IDT. AICS-0097-141 ACTN2-mEGFP MYH7WT/G256E iPSCs were dissociated into single-cell suspension using Accutase. Transfections were performed using the Lonza 4D Nucleofector Nucleofection Transfection System. 1.6μl 61μM Alt-R Cas9 was incubated for 10 min at room temperature with

2.4 µl of 50µM sgRNA, precomplexed and co-transfected with 1 µl of 100 µM ssODN donor. 5 × $10^5$ cells were resuspended in 20µl nucleofection buffer and added to the 5 µl of ssODN, Cas9 protein, and sgRNA. The electroporation program used was CA-137. Cells were then immediately plated onto Matrigel–coated six-well plate with mTeSR1 medium supplemented with 10 µM ROCK inhibitor. Transfected cells were cultured as described under iPSC Cell culture above for 3–4 days until the transfected culture had recovered to ~70% confluence. Cells were subsequently passaged very sparsely, and single-cell clones were picked. Clones were sequenced via Sanger sequencing and karyotyped to confirm successful gene editing.

*Collection and lysis of hiPSC-CMs for global proteome analysis*

Frozen day 12 hiPSC-CMs containing either wild type MYH7 alleles (WT), a homozygous MYH7 G256E mutation (HOM) or a heterozygous MYH7 G256E mutation (HET) were thawed and 2M CMs were plated per 6 well. For global proteome analysis, 2 wells of a 6-well-plate were combined for each sample. In total, 2 clones of WT hiPSC-CMs (clones 113 and 174), 2 clones of HOM hiPSC-CMs (clones 15 and 17) and 2 clones of HET hi-PSC-CMs (clones 102 and 141) were analysed in biological triplicates. Cells were washed twice with ice-cold PBS, before addition of 200 µL lysis buffer (6 M Guanidine, 100 mM Tris-HCl, pH 8.5, 95°C) to each 6-well and collection using a cell scraper. Cell lysate was transferred to a 1.5 mL protein loBind reaction tube, incubated at 95°C for 10 min (Eppendorf Thermoshaker), sonicated for 3 cycles of 30 sec on / 30 sec off at 4°C (Bioruptor) and stored frozen.

*Protein digestion and peptide desalting*

The sample preparation was performed as described previously with minor modifications.[7] In short, samples were centrifuged for 10 min at 14000 rpm, the supernatant was transferred to a new protein loBind reaction tube and sonicated for 10 min (Branson 2800). Protein concentration was determined using a Pierce™ BCA Protein Assay Kit (Thermo Scientific). Further, 150 µg protein of each sample was reduced and alkylated at a final concentration of 5 mM tris (2-carboxyethil) phosphine (TCEP, Sigma) and 10 mM chloroacetamide (CAA, Sigma) for 10 min at 95°C protected from light. Samples were then diluted to 2 M guanidine-HCl using 50 mM Tris pH 8.5 and digested by 1:50 (w/w) Lys-C (Sequencing grade, Wako) for 1 h at 25°C and 700 rpm. Proteins were further diluted to 0.5 M guanidine-HCl with 50 mM Tris pH 8.5 and digested with trypsin (Sequencing grade, Promega) 1:100 (w/w) over night at 37°C and 700 rpm. Digestion was quenched by addition of trifluoroacetic acid (TFA) to a final concentration of 0.5% (pH ~2). Digested peptides were desalted on a solid phase extraction system (C18 SepPak, 1 cc, 50 mg, Waters Corp.; wash solvent 0.1% TFA and 0.1% FA). Peptides were eluted stepwise with 40% acetonitrile (ACN) and 60% ACN and dried by vacuum centrifugation (Eppendorf, Concentrator Plus).

*Tandem-mass-tag (TMT) labeling*

TMT labelling was performed using a standard protocol, as described by Zecha and colleagues.[8] Dried peptides were resuspended in 22 µL 100 mM HEPES pH 8.5 and peptide concentration was determined (Lunatic, Unchained labs). 40 µg peptides of each sample were labeled with 100 µg of tandem-mass tag (TMT) reagents resuspended in 5 µL 100% anhydrous ACN. The final reaction conditions were as follows: TMT-to-peptide ratio 2.5:1, 1.6 µg/µL peptide concentration, 20% ACN, 80 mM HEPES pH 8.5. The TMT-peptide mixture was incubated for 1 h at 25°C and 750 rpm, and the labeling reaction was quenched by addition of 5% hydroxylamine to a final concentration of 1% for 15 min at 25°C and 750 rpm. Samples were combined, concentrated for 25 min to volatilize ACN (Eppendorf Concentrator Plus), acidified to pH 2 by stepwise addition of 10% TFA, and desalted as described above.

*Fractionation at high pH*

Dried peptides were resuspended in 10 mM triethylammonium bicarbonate (TEAB), and the final peptide concentration was determined (Lunatic, Unchained labs). 20 µg TMT-labeled peptides (concentration 1 µg/µL) were fractionated using a Kinetex C18 column (100 Å, 2.6 µm, 0.3 mm x 150 mm, Phenomenex) on an EASY-nLC 1200 system (Thermo Scientific) operating at a flow rate of 2 µL/min with two buffer lines (Buffer A: 10 mM TEAB, Buffer B: 80% ACN, 10 mM TEAB). Peptides were separated over a 100 min gradient (3-40% B in 57 min, 40-60% B in 5 min, 60-95% B in 10 min, 95% B for 10 min, 95-3% B in 10 min, 3% B for 8 min) and 24 concatenated fractions were collected at 30 s sampling intervals in 0.1% FA (twin.tec® PCR Plate 96 loBind, skirted, Eppendorf). The resulting fractions were dried down by vacuum centrifugation and resuspended at a concentration of 0.1 µg/µL in 2% ACN, 0.1% TFA prior to LC-MS/MS analysis.

*LC-MS/MS analysis*

The resulting 24 fractions of TMT-labelled peptides were analyzed on an Orbitrap Ascend Tribrid mass spectrometer coupled to a Vanquish Neo UPHLC system (Thermo Fisher Scientific). Peptides were separated on a 25 cm x 75 µm Aurora Gen2 column (IonOpticks) at a flow rate of 400 nL/min over a 101 min multi-step linear gradient (Buffer A: 0.1% formic acid, Buffer B: 0.1 % formic acid, 80% ACN; 1-5% B in 2 min, 5-17% B in 55 min, 17-25% B in 21 min, 25-35% B in 13 min, 35-85% B in 3 min, 85% B for 7 min). Column effluent was directly ionized in a nano-electrospray ionization source operated in positive ionization mode and electrosprayed into the mass spectrometer. Full scan mass spectra were acquired in the orbitrap at a resolution of 60,000, a scan range of 400-1400 m/z and a normalized AGC target of 100 % or a maximum injection time of 123 ms. Most intense precursor ions (intensity $\geq 2.5 \times 10^4$) with charge states 2-6 and monoisotopic peak determination set to 'peptide' were selected for MS/MS fragmentation by higher-energy collisional dissociation (HCD) at 35 % collision energy in a data dependent mode. The MS2 isolation width was set to 0.7 m/z and the duration for dynamic exclusion was set to 60s. MS/MS spectra of fragment ions were analyzed in the orbitrap at a resolution of 45,000 and a scan range of 110-2000 m/z (AGC target 200 %, maximum fill time of 91 ms). All data were acquired with the FAIMS Pro Duo™ Interface at -45 CV and the cycle time was fixed to 2.4s.

*Proteomics Raw data processing*

Thermo raw files were analyzed with FragPipe v21.1 (MSFragger v4.0, IonQuant v1.10.12, Philosopher v5.1.0). In short, the TMT16 workflow was used with default MSFragger settings, including oxidation of methionine, acetylation of the protein N-terminus, and N-terminal TMTpro labeling as variable modification. Fixed modifications included carbamidomethylation of cysteine and TMTpro labeling of lysine. The data was first searched against a human canonical protein sequence database (SwissProt, downloaded on 06/03/2024), supplemented with common contaminants, a full-length sequence of the MYH7_G256E mutant, and decoy sequences. The MYH7_G256E entry was then modified to contain only the identified peptide spanning the amino acid substitution (residues 250-257), and this FASTA file was used in the final search. Isobaric labeling-based quantification was performed with the label type set to TMT-18. Quantification results were filtered for 'No PSMs mapping to multiple genes in dataset'. MS1 intensities and top 3 ions were disabled for the conversion of ratios to abundances. Finally, intensities were log2 transformed and median centered prior to downstream analysis. The mass spectrometry proteomics data have been deposited to the ProteomeXchange Consortium via the PRIDE[9] partner repository with the dataset identifier PXD054863.

*Proteomics Statistical analysis*

Normalized quantitative reports were imported into RStudio (v2023.09.1). Peptide-level quantification was utilized to assess the ratio of the mutant MYH7_G256E peptide across groups, while gene-level quantification was employed for all other analyses. Differential expression analysis was performed using the 'dream' function from the varianceParition package (v.1.32.5).[10] A linear mixed model was fitted for each gene separately, with 'Genotype' specified as a fixed effect and 'Clone' treated as a random effect to account for within-clone correlation. The 'eBayes' function from the limma package (v3.56.2)[11], utilizing the limma-trend method, was employed to compute moderated t-statistics through empirical Bayes moderation of the standard errors. The results for the contrast of interest were summarized using the 'toptable' function while adjusting p-values using the Benjamini-Hochberg procedure to control the false discovery rate. All visualizations were generated using the ggplot2 package (v3.4.4). Functional enrichment was performed using the Database for Annotation, Visualization, and Integrated Discovery (DAVID). Upregulated and downregulated DEPs were submitted to DAVID Bioinformatics Resources Database (https://david.ncifcrf.gov/home.jsp) for identifying enriched GO-biological process and GO-molecular function terms. Functional annotation clustering was then used to identify functional groups within the identified terms.

### 2. SI Figure Legends

**Suppl. Fig. 1:** (A) Feature plot of H251N CMs showing gene expression of CM markers *TNNT2* (I) and *MYH7* (II), endothelial marker *ACE* (III) and fibroblast marker *DCN* (IV). (B) Feature plot of G256E CMs showing gene expression for the same CM, endothelial and fibroblast markers (I-IV)

**Suppl. Fig. 2:** (A) Heatmap showing expression of top 50 DEGs between H251N HET CMs and WT CMs, sorted by clone and experimental run (WT n=612, HET n=846). (B) Volcano plot showing all DEGs between H251N HET CMs and WT CMs. Grey color indicating no significant regulation (NS), red color indicating statistically significant regulation above foldchange cutoff > 0.5. Green and Blue color indicating regulation either only above the significance or only above fold change threshold. Significance determined by Seurat's two-sided Wilcoxon rank-sum test with Bonferroni correction for multiple testing. (C) Top GO-term enrichment clusters for identified DEGs.

**Suppl. Fig. 3:** (A) Heatmap showing expression of top 28 DEGs between G256E HET CMs and WT CMs, sorted by clone and experimental run (WT n=1332, HET n=1106) (B) Volcano plot showing all DEGs between G256E HET CMs and WT CMs. Grey color indicating no significant regulation (NS), red color indicating statistically significant regulation above foldchange cutoff > 0.5. Green and Blue color indicating regulation either only above the significance or only above fold change threshold. Significance determined by Seurat's two-sided Wilcoxon rank-sum test with Bonferroni correction for multiple testing.

**Suppl. Fig. 4:** (A) UMAP distribution of H251N and WT CMs split by experimental run and clone. (B) UMAP distribution of G256E and WT CMs split by experimental run and clone. (C) Run-to-run DEG expression difference split by clone. DEGs were defined as log fold change >0.5 & adjusted p-value < 0.05. Each dot represents the average run-to-run expression difference of one DEG for a specific clone. N(H251N clones) = 165; n(G256E clones) = 28 DEGs. Salmon color indicating WT clone and blue color indicating mutant clone. Data are presented as mean ± SEM.

**Suppl. Fig. 5:** Raw Digital-Droplet PCR readouts for WT-oligo and mutant-oligo intensity for each G256E and WT CM clone across three biological replicates for each clone.

**Suppl. Fig. 6:** (A) Schematic of experimental set-up. (B) Distribution of *MYH7* G256E mutant allele expression fraction per cell (MUT / [WT + MUT]) for two independent sequencing runs (Run 1 [n=23] and Run 2 [n=46]), including only cells with ≥5 reads per cell. (C) UMAP visualization of single-cell transcriptomic data showing cluster assignments and expression of cardiac marker genes (*TNNT2* and *MYH7*), along with quality control metrics (nFeature_RNA and nCount_RNA). (D) Correlation between mutant allele fraction and hypertrophy-associated module score (NPPB, APOE, ANKRD1) across single cells. Correlation between allelic fraction and module score was assessed using a two-sided Pearson correlation test.

**Suppl. Fig. 7:** (A) Representative images of WT, G256E HET and H251N HET CMs showing, from left to right, α-actinin–GFP fluorescence, sarcomere orientation (color indicates local sarcomere orientation relative to the horizontal axis [cyan = horizontal, red = ±90°, vertical]), and sarcomere tracking (color-coded by Z-to-Z distance from long [pink] to short [blue]). Scale bars: 50 µm (field of view, sarcomere orientation), 10 µm (sarcomere tracking). (B) Quantification of contraction velocity (CV), fractional sarcomere shortening (FSS), sarcomere orientation (SO), and relaxed Z-to-Z distance (RZ) by clone in G256E HET, H251N HET and WT cardiomyocytes (CV n = 15; FSS, SO, RZ n = 10).

**Suppl. Fig. 8:** (A) Feature plot of G256E CMs after isoproterenol treatment showing gene expression of CM markers *MYH7* (I) and *TNNT2* (II), fibroblast marker *DCN* (III) and endothelial marker *ACE* (IV). (B) UMAP distribution of G256E and WT CMs after isoproterenol split by experimental run and clone. (C) Quantification of run-to-run expression variability of DEGs between independent scRNAseq runs. DEGs were defined as log fold change >0.5 & adjusted p-value < 0.05. Each dot represents the average run-to-run expression difference of one DEG for a specific clone. N(G256E) = 140 (5 clones, 28 DEGs); n(G256E + Iso) = 245 (5 clones, 49 DEGs). (D) Quantification of clone-to-clone expression variability of DEGs between three isogenic G256E HET CM clones and two isogenic WT CM clones with and without isoproterenol treatment. DEGs were defined as log fold change >0.5 & adjusted p-value < 0.05. Each dot represents the average clone-to-clone expression difference of one DEG. N(WT & HET) = 28, n(WT +Iso & HET +Iso) = 49. Data are presented as mean ± SEM Statistical significance was determined by by linear mixed-effects models (lmerTest) with genotype as a fixed effect and clone as a random effect (C) and ordinary one-way ANOVA with post-hoc Tukey's multiple comparisons test (D).

**Suppl. Fig. 9:** (A) Heatmap showing expression of top 49 DEGs between G256E HET and WT CMs after isoproterenol treatment, sorted by clone and experimental run (WT n=1604, HET n=5958) (B) Volcano plot showing all DEGs between G256E HET and WT CMs after isoproterenol treatment. Grey color indicating no significant regulation (NS), red color indicating statistically significant regulation above foldchange cutoff > 0.5. Green and Blue color indicating regulation either only above the significance or only above fold change threshold. P-values determined by Seurat's two-sided Wilcoxon rank-sum test with Bonferroni correction for multiple testing. (C) Top GO-term enrichment clusters for identified DEGs.

**Suppl. Fig. 10:** (A-C) Quantification of mitochondrial (A), glycolytic (B) and total (C) ATP production rate in WT vs G256E CMs split by individual clones with and without isoproterenol treatment. Measured by ATP Rate Assay. (WT C113: basal n=3, iso n=4; WT C174: basal n=4, iso n=4; HET C102: basal n=4, iso n=4; HET C141: basal n=3, iso n=3; HET C157: basal n=4, iso n=3). Data are presented as mean ± SEM. Statistical significance was determined by unpaired t test. *$P < 0.05$, **$P < 0.01$, ****$P < 0.0001$.

**Suppl. Fig. 11:** (A) Feature plot of G256E CMs after isoproterenol treatment showing gene expression of CM markers *MYH7* (I) and *TNNT2* (II), fibroblast marker *DCN* (III) and endothelial marker *ACE* (IV). (B) UMAP distribution of G256E and WT CMs after isoproterenol split by experimental run and clone.

**Suppl. Fig. 12:** (A) Heatmap showing expression of top 50 DEGs between G256E HOM and WT CMs, sorted by clone and experimental run (WT n=1304, HOM n=980) (B) Volcano plot showing all DEGs between G256E HOM and WT CMs. Grey color indicating no significant regulation (NS), red color indicating statistically significant regulation above foldchange cutoff > 0.5. Green and Blue color indicating regulation either only above the significance or only above fold change threshold. P-values determined by Seurat's two-sided Wilcoxon rank-sum test with Bonferroni correction for multiple testing. (C) Top GO-term enrichment clusters for identified DEGs.

**Suppl. Fig. 13:** (*A)* Total MYH7 protein abundance and MYH7 G256E peptide abundance in each sample. (*B)* Heatmap displaying hierarchical clustering of protein expression profiles across different genotypes and clones. (C) Correlation of G256E HET vs WT fold changes in proteomic and transcriptomic dataset. Black color indicating positive gene/protein expression correlation & grey color indicating negative gene/protein expression correlation.

**Suppl. Fig. 14:** (A) Dotplot showcasing expression of marker genes consistently regulated across all *MYH7* G256E experimental conditions.

**Suppl. Fig. 15:** (A) Boxplots of protein abundance of ANKRD1 and NPPB. Data presented using boxplots show the median and interquartile range (IQR). Fc = log2 fold change. P-values were calculated using moderated t-statistics computed through empirical Bayes moderation of the standard errors, as implemented in the limma package. P-values were further adjusted for multiple comparisons using the Benjamini-Hochberg procedure to control the false discovery rate. Two clones of WT hiPSC-CMs (clones 113 and 174), 2 clones of HOM hiPSC-CMs (clones 15 and 17) and 2 clones of HET hi-PSC-CMs (clones 102 and 141) were analysed in biological triplicates. (B) Quantification of Western Blot Gels measuring abundance of ANKRD1 and NPPB relative to TNNT2 in WT and *MYH7* G256E HET and HOM CMs. Two clones of WT hiPSC-CMs (clones 113 and 174), 2 clones of HOM hiPSC-CMs (clones 15 and 17) and 3 clones of HET hiPSC-CMs (clones 102,141 and 157) were analysed. (C) Corresponding Western Blot Gels.

**Suppl. Fig. 16:** Heatmaps showing cluster-specific expression patterns across day 30 CM datasets for G256E HET CMs, H251N HET CMs, G256E HET CMs + isoproterenol, and G256E HOM CMs. Cells are grouped by transcriptional clusters, with selected marker genes shown on the y-axis.

**Suppl. Fig. 17:** (A) UMAP representation of maturation media treated (six days) G256E HET CMs and the isogenic WT counterpart clustered by the presence of the G256E mutation. Unbiased clustering of G256E HET and WT hiPSC-CMs. Relative contribution to each cluster of WT and G256E HET hiPSC-CMs. Comparison of *TNNT2* gene expression in WT and G256E hiPSC-CMs. (B) Dotplot of *MYL2* and *MYL7* expression across WT and G256E HET CMs cultured in standard versus maturation media. (C) Gene expression of *NPPB* and *ANKRD1* in WT and G256E HET CMs with and without maturation media treatment. Statistical significance was determined by a two-sided Wilcoxon rank-sum test with Bonferroni correction as implemented in Seurat (C).

**Suppl. Fig. 18:** (A) UMAP representation of day 60 G256E HET CMs and the isogenic WT counterpart clustered by the presence of the G256E mutation. Unbiased clustering of G256E HET and WT hiPSC-CMs. Relative contribution to each cluster of WT and G256E HET hiPSC-CMs. Comparison of *TNNT2* gene expression in WT and G256E hiPSC-CMs. (B) Dotplot of *MYL2* and *MYL7* expression across WT and G256E HET CMs in day 30 versus day 60 conditions. (C) Gene expression of *NPPB* and *ANKRD1* in WT and G256E HET CMs in day 30 and day 60 conditions. Statistical significance was determined by a two-sided Wilcoxon rank-sum test with Bonferroni correction as implemented in Seurat (C).

**Suppl. Table 1:** Identified DEGs between H251N HET and WT CMs

**Suppl. Table 2:** Identified DEGs between G256E HET and WT CMs

**Suppl. Table 3:** Identified DEGs between G256E HET and WT CMs after isoproterenol treatment

**Suppl. Table 4:** Identified DEGs between G256E HOM and WT CMs

**Suppl. Table 5:** Identified DEPs between G256E HET and WT CMs

**Suppl. Table 6:** Identified DEPs between G256E HOM and WT CMs

**Suppl. Table 7:** List of DEGs specific to only G256E HOM and H251N HET CMs, only H251N HET CMs and only G256E HOM CMs

P-values in suppl. Tables were determined by Seurat's two-sided Wilcoxon rank-sum test with Bonferroni correction for multiple testing.
